# Enzymatic RNA Biotinylation for Affinity Purification and Identification of RNA-protein Interactions

**DOI:** 10.1101/2020.05.31.122085

**Authors:** Kayla N. Busby, Amitkumar Fulzele, Dongyang Zhang, Eric J. Bennett, Neal K. Devaraj

## Abstract

Throughout their cellular lifetime, RNA transcripts are bound to proteins, playing crucial roles in RNA metabolism, trafficking, and function. Despite the importance of these interactions, identifying the proteins that interact with an RNA of interest in mammalian cells represents a major challenge in RNA biology. Leveraging the ability to site-specifically and covalently label an RNA of interest using *E. Coli* tRNA guanine transglycosylase and an unnatural nucleobase substrate, we establish the identification of RNA-protein interactions and the selective enrichment of cellular RNA in mammalian systems. We demonstrate the utility of this approach through the identification of known binding partners of 7SK snRNA via mass spectrometry. Through a minimal 4-nucleotide mutation of the long noncoding RNA HOTAIR, enzymatic biotinylation enables identification putative HOTAIR binding partners in MCF7 breast cancer cells that suggest new potential pathways for oncogenic function. Furthermore, using RNA sequencing and qPCR, we establish that an engineered enzyme variant achieves high levels of labeling selectivity against the human transcriptome allowing for 145-fold enrichment of cellular RNA directly from mammalian cell lysates. The flexibility and breadth of this approach suggests that this system could be routinely applied to the functional characterization of RNA, greatly expanding the toolbox available for studying mammalian RNA biology.

## INTRODUCTION

Proper regulation of RNA metabolism, with elaborate networks of RNAs and their associated RNA-binding proteins (RBPs), underlies cellular homeostasis. Post-transcriptional gene regulation is dictated by the interactions of coding and noncoding RNAs with RNA-binding proteins to ensure the accurate coordination of RNA processing, transport, translation, and degradation. For example, the 3′ untranslated regions (UTRs) of mRNAs are known to be regulatory hubs for RNA-binding proteins that control mRNA expression level and stability.^1^ Furthermore, long non-coding RNAs, through their interactions with proteins, have been shown to be involved in various regulatory processes such as chromatin remodeling and transcriptional activation.^2^ These networks of RNA-protein interactions have been implicated in numerous human diseases, ranging from cancer to neurodegeneration.^3,4^ Therefore, the characterization of these RNA-protein interactions is crucial to our understanding of RNA biology and the development of innovative RNA-centric therapeutic strategies.

With increasing appreciation for the role of RNA-protein interactions in gene regulation and disease, there is a growing need for methods enabling their identification via protein-centric and RNA-centric approaches. Protein-centric approaches, which allow for identification of RNA transcripts bound to a given protein, have progressed rapidly in the past decade as next-generation sequencing technologies have become more widely available. Techniques such as RNA immunoprecipitation (RIP-seq) and crosslinking and immunoprecipitation (CLIP-seq) are widely used.^5^ While conceptually simple, RNA-centric approaches that allow for identification of proteins bound to a given RNA transcript are not as common as protein-centric approaches for two related reasons: First, a typical mRNA is about 3 orders of magnitude less abundant than a typical protein in mammalian cells,^6^ and second, mass spectrometry is needed to identify cognate proteins, which, unlike sequencing, is not amenable to signal amplification. Because of these challenges, RNA affinity purification and cognate protein identification remains a challenge in RNA biology. Despite these obstacles, RNA-centric studies are essential to study the function and regulation of targeted RNA transcripts.

Existing noncovalent approaches for RNA-centric interaction mapping include biotinylated antisense oligonucleotides (ASOs) and naturally occurring RNA-protein partners, such as MS2 tagging. Because cellular RNA exhibits intricate secondary and tertiary structure and is typically coated with various RNA binding proteins, it has historically been challenging to design antisense probes that efficiently purify an RNA of interest. The use of RNA hybridizing antisense probes to selectively purify the long-noncoding RNA *Xist* from mammalian cells after cell crosslinking was recently achieved through two similar approaches, RAP-MS and CHIRP-MS.^7–9^ To attain adequate selectivity and purification efficiency, both strategies utilized biotinylated antisense probes that tile the length of the 18-kb RNA. While this work represents an important advance in the field, it is unclear how general these methods are for long noncoding RNAs (lncRNAs) or mRNAs with different sequences, splice variants, lower expression levels, and a more typical length (∼1-2 kb).^10,11^ Antisense oligonucleotides have also been demonstrated using a two-step purification of mRNA transcripts with a preliminary polyA purification, in addition to the analysis of ribosomal RNA.^12,13^ While these approaches extend the utility of antisense oligonucleotides, ribosomal RNA is extremely abundant, and the two-step approach relies on polyA purification, and thus is not applicable to nonpolyadenylated RNAs. Furthermore, optimization of ASO sequence and consideration of RNA base pairing availability is required. In addition to ASO approaches, naturally occurring RNA-protein interactions, such as the interaction between MS2 coat protein and its cognate RNA hairpin, have been developed as a method for RNA affinity purification.^14–16^ To facilitate affinity purification, several MS2 cognate hairpins are appended to the RNA of interest, and the MS2 coat protein is expressed as a fusion with another affinity protein. However, limited by the relatively low affinity of MS2 protein with the target RNA (∼3 nM), the performance of such systems is highly dependent on the expression level of the RNA of interest.^17^ Thus, a stronger affinity handle is ideal in studying RNA-protein interactions, especially for less abundant RNA species. For instance, the affinity between biotin and streptavidin is ∼1 fM, making biotin a excellent affinity handle to tag an RNA of interest and capture its interactome.^18^

Enzymatic covalent biotinylation of RNA has various potential advantages over noncovalent approaches for affinity purification. First, due to the covalent interaction, it would have the potential to facilitate recovery of less abundant RNA transcripts. Furthermore, by providing a robust covalent linkage, stringent purification conditions of the RNA-protein complexes can be utilized, making such purifications much more accessible. Common approaches for biotinylation of RNA transcripts include *in vitro transcription* with biotinylated precursors, or ligation of a biotin to the end of an RNA transcript.^19,20^ However, these existing methods for covalent biotinylation of RNA do not target a specific RNA transcript, and would therefore not be used for the purification of a target cellular RNA. While enzymatic methods for RNA modification have been developed, many of these methods are not selective for a target transcript, and none of these methods have been demonstrated to achieve selective covalent labeling of a target RNA against the transcriptome.^21^ Few examples exist whereby enzymes can be utilized to have been harnessed to accomplish site-specific covalent modification of RNA.^21–24^ In previous approaches, reactive precursors such as primary amines and common metabolites were utilized that are likely not orthogonal to mammalian systems.^25^ Additionally, these approaches have required large stretches of RNA sequence, such as an entire tRNA structure, for recognition and modification.

Recently, we have shown that a bacterial RNA modifying enzyme, tRNA guanine transglycosylase (TGT), can be utilized to covalently label an RNA of interest bearing a short, encoded hairpin recognition motif.^26,27^ Based on the minimal recognition motif, direct covalent modification, and potential for selectivity in mammalian systems, we sought to apply this methodology, called RNA Transglycosylation at Guanosine (RNA-TAG) to the study of RNA-protein complexes in mammalian cells (Figure 1). Herein, we demonstrate the use of RNA-TAG as a novel and effective approach to site-specifically biotinylate and isolate RNAs for the analysis of RNA-protein interactions using immunoblotting and quantitative mass spectrometry. We also demonstrate that native RNA structures can be transformed into enzymatic labeling sites in order to minimize perturbation of the native RNA sequence. Furthermore, while previous imaging studies have suggested that TGT has some degree of selectivity against mammalian transcriptomes,^26,28^ we demonstrate that the use of dimerized TGT enables efficient and substantially more selective labeling and affinity purification of an expressed RNA of interest within cell lysates, demonstrating the potential breadth of this approach to include the study of cellular RNA interactions.

**Figure 1.**
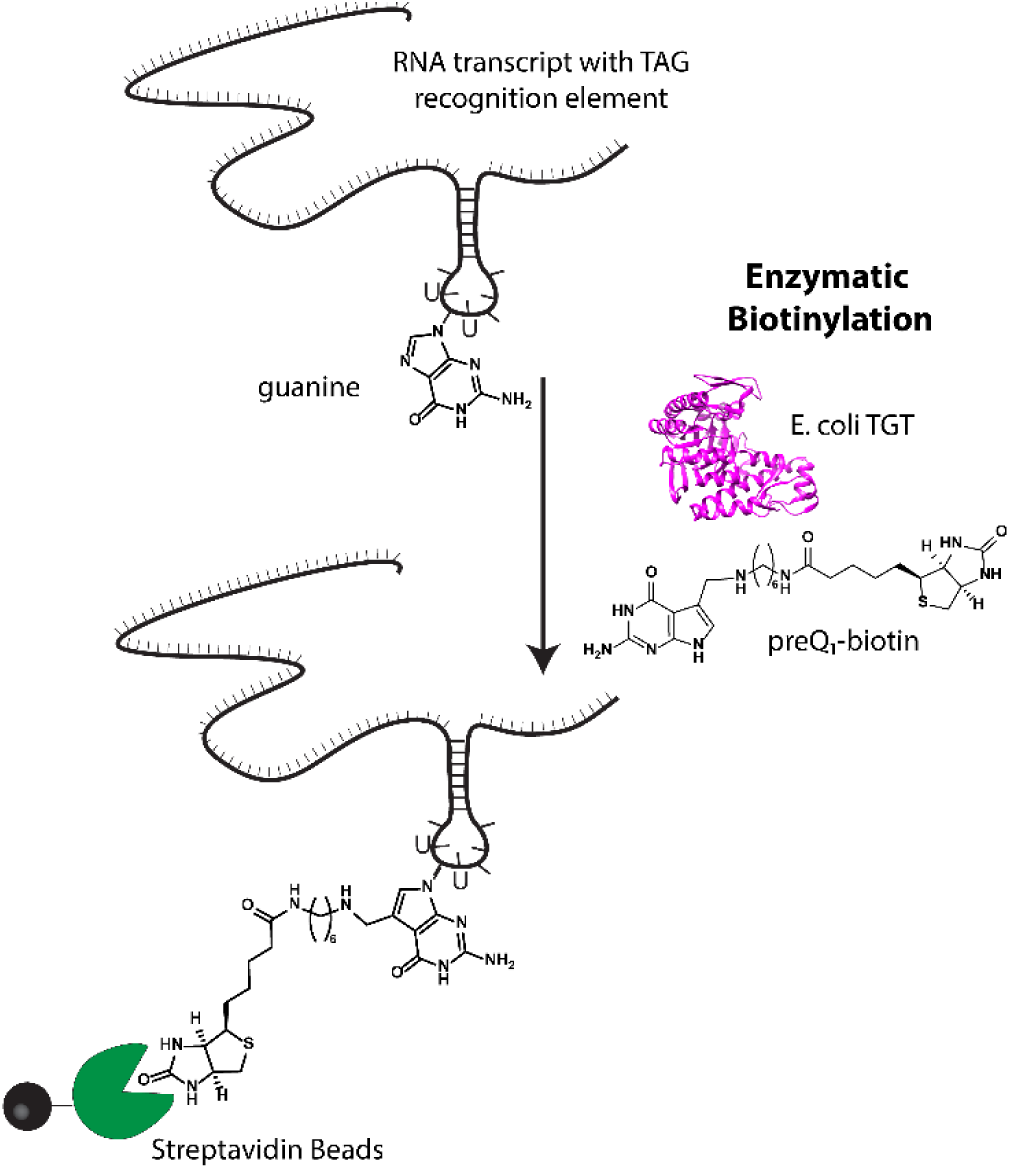
Enzymatic RNA labeling with RNA-TAG. Using bacterial tRNA Guanine Transglycosylase (TGT), an RNA of interest containing a TGT recognition hairpin is labeled with a derivative of the natural substrate, preQ_1_.

## RESULTS AND DISCUSSION

### Detection of RNA-Protein Interactions by Immunoblot

We first sought to apply RNA-TAG to study the protein binding partners of human mRNA targets. As a proof-of-concept, we selected a fragment of HDAC2 mRNA known to bind the important RNA binding protein HuR,^29^ as well as full-length β-actin mRNA, which is also a known binding partner to HuR.^30,31^ We designed RNA constructs for each of these sequences, termed HDAC2-TAG and β-actin-TAG, which had an appended 25-nucleotide TGT recognition hairpin at the 3’ end surrounded by a short, unstructured linker at either side. Furthermore, we designed an RNA construct, termed Control-TAG, that encoded the TGT recognition hairpin and linker regions, but did not have a specific RNA target encoded (Figure 2A, B). To test whether RNA transcripts biotinylated using RNA-TAG could be utilized for enrichment of cognate RNA binding proteins, we first labeled each transcript through treatment with *E. coli* TGT and a preQ_1_-biotin conjugate. The resulting purified RNA product was then folded and incubated directly with cellular lysates prepared from HeLa cells in order to allow binding of the labeled RNA with cognate RNA binding proteins. Streptavidin-mediated affinity purification was used to enrich the target RNA-protein complexes, which were then analyzed via immunoblot (Figure 2C). Gratifyingly, we found that HuR was enriched in the HDAC2-TAG and β-actin-TAG samples, but not in the Control-TAG sample. Furthermore, β-tubulin, which is not expected to interact with these RNAs, was not enriched in any samples, indicating specificity of the enrichment. One potential advantage of this approach is the strength of the streptavidin-biotin interaction, potentially allowing for RNA-protein complexes to be detected at extremely low concentrations. In order to examine this, we carried out RNA-protein complexation, affinity purification, and immunoblotting with decreasing amounts of labeled HDAC2-TAG RNA (Figure 2D). The HuR binding partner was detected with as little as 1.5 pg of biotinylated RNA, corresponding to a concentration in the sub-picomolar range (10^−13^ M). For comparison, an average mRNA transcript has approximately 20 copies per cell, corresponding to concentrations in the picomolar range (10^−11^ M), based on a cell volume of 2.6 pL.^6,32^ We also tested whether RNA biotinylation could be carried out at lower concentrations of RNA, and found that RNA was labeled with good efficiencies at concentrations down to 1 pM (Supplementary Figure S1).

**Figure 2.**
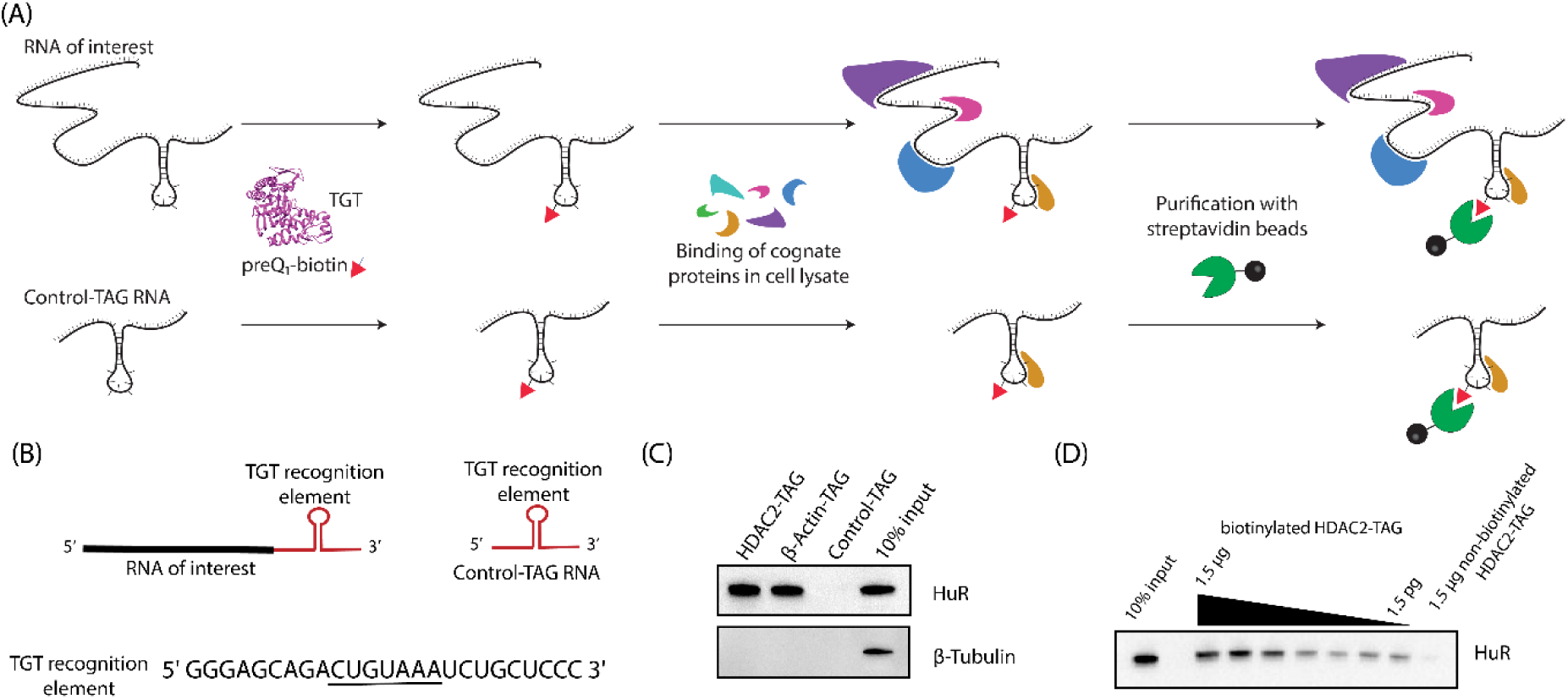
Analysis of RNA-protein interactions using RNA-TAG. (A) An RNA of interest with an appended TAG sequence is biotinylated via treatment with *E. Coli* TGT and preQ_1_-biotin and allowed to bind cognate RNA-binding proteins, alongside a control RNA for analysis of bound proteins (B) RNA constructs used for western blotting experiments contain an RNA of interest (HDAC2-HuR binding domain or β-Actin mRNA) followed by a 25-nt TGT recognition element with the indicated sequence, where the loop region is underlined, surrounded by short spacer regions. Control-TAG RNA has no RNA of interest inserted. (C) RNAs biotinylated with RNA-TAG were used directly to analyze cognate proteins via western blotting by incubation of 1 pmol labeled RNA with 40 μg cell lysate. (D)HDAC2-TAG-HuR complex can be detected using reduced RNA concentrations, as shown by a titration of RNA from 1.5 μg to 1.5pg.

### Identification of RNA-protein interactions by quantitative proteomics

Encouraged by our immunoblot data, we sought to determine whether this methodology for the purification of enzymatically biotinylated RNA-protein complexes could be applied to the discovery of RNA-protein interactions using quantitative mass spectrometry (Figure 3A). In addition to the model RNA HDAC2-TAG, we used 7SK as a target for proteomics experiments. 7SK is an abundant small nuclear RNA (snRNA) that is approximately 330 nucleotides in length and is involved in transcription regulation. 7SK snRNA has been extensively studied due to its role in the regulation of the positive transcription elongation factor b (P-TEFb).^33^ This activity is regulated through its interaction with proteins such as lupus antigen-related protein 7 (LARP7), methylphosphate capping enzyme (MePCE), and HMBA-induced proteins 1 and 2 (HEXIM1/2). Proteins interacting with 7SK snRNA have recently been reported using the CHIRP-MS method,^8,34^ which utilizes tiling biotinylated antisense probes for RNA affinity purification; therefore, 7SK is an ideal target for validation of RNA-TAG affinity purification. In order to mediate RNA-TAG labeling of 7SK, we encoded a TGT recognition element on the 5’ end (7SK-TAG, Figure 3B).

**Figure 3.**
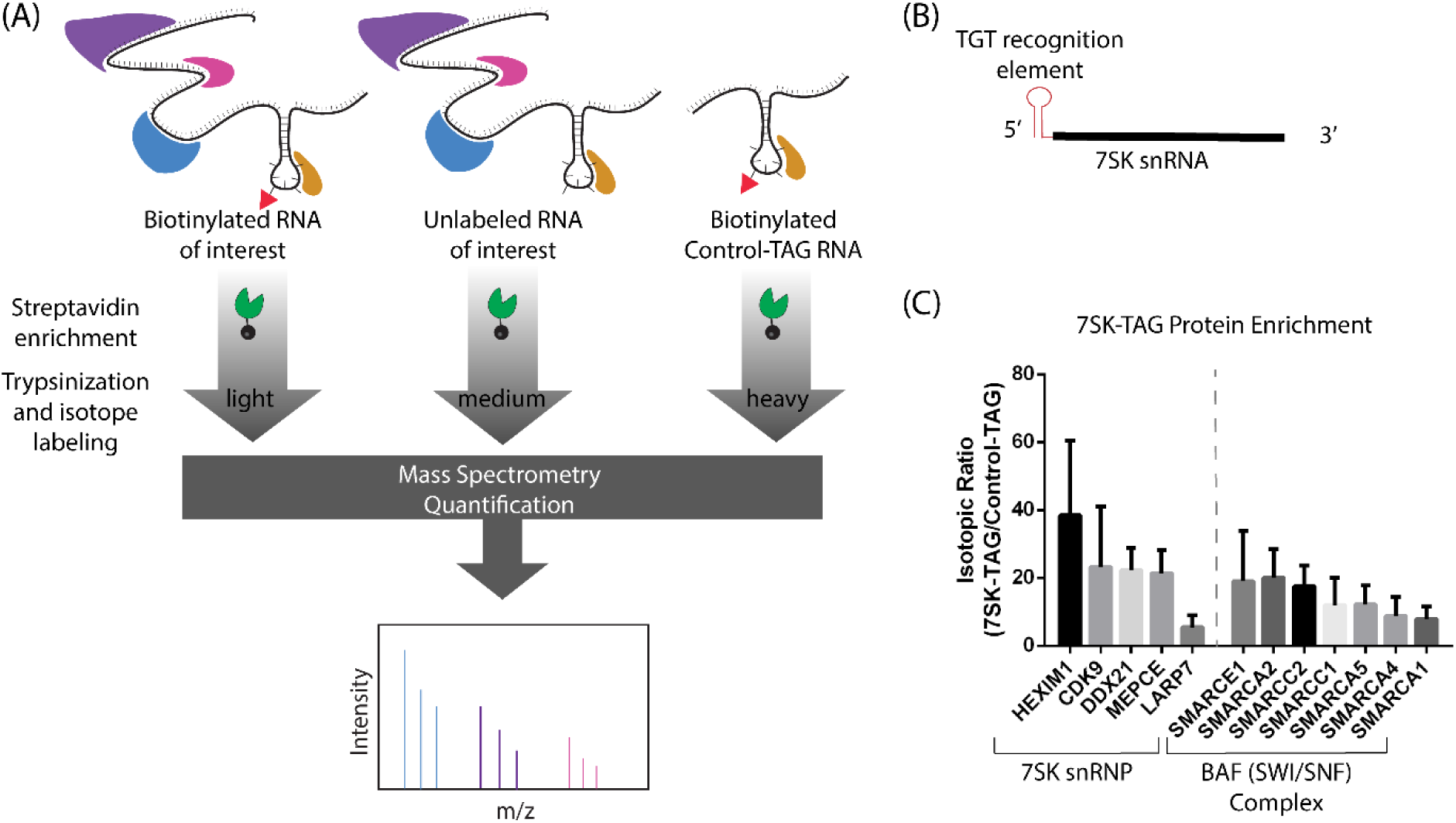
Quantitative proteomics to identify RNA-protein interactions. (A)The affinity purification of the biotinylated RNA of interest was carried out alongside two controls, an unlabeled RNA of interest and a biotinylated Control-TAG transcript. Isotope labeling was carried out using reductive dimethylation, samples were pooled and analyzed with LCMS/MS. (B) 7SK-TAG RNA construct with TAG sequence appended directly to the 5’ end (C) Isotopic ratios (biotinylated 7SK-TAG/biotinylated Control-TAG) for enriched proteins that are known interactors with 7SK snRNA or part of the BAF complex known to interact with 7SK snRNP (mean ± S.D.) Number of replicates were as follows: n=5 (CDK9, DDX21, MEPCE, LARP7, SMARCA5), n=4 (SMARCE1, SMARCA2, SMARCC1), n=3 (HEXIM1, SMARCA4, SMARCA1, SMARCC2)

In order to ascertain which proteins were enriched by binding these target RNAs, we carried out quantitative mass spectrometry using a reductive demethylation-peptide labeling strategy.^35^ To control for background proteins and nonspecific proteins, three samples were compared for each target RNA, consisting of biotinylated target RNA (HDAC2-TAG or 7SK-TAG), unlabeled target RNA (HDAC2-TAG or 7SK-TAG), and biotinylated Control-TAG RNA. As before, biotinylation of HDAC2-TAG, 7SK-TAG, and Control-TAG was achieved through treatment with preQ_1_-biotin and *E. Coli* TGT. Each RNA sample was incubated with cell lysate to allow cognate binding proteins to bind, and complexes were affinity purified, with five biological replicates for each target. In order to verify the biotinylation of the RNA constructs, Northern blots were performed and transferred RNA was developed directly using Streptavidin-HRP (Supplementary Figure S2). Digested and labeled peptides were pooled and analyzed by liquid chromatography-mass spectrometry. Ratios between the experimental samples and the two control samples were calculated and used to identify highly enriched proteins. To generate lists of interacting proteins, the experimental sample (biotinylated target RNA) was compared to both controls (unlabeled target RNA and biotinylated control-TAG RNA). Proteins detected in at least three replicates with an average enrichment greater than 3-fold against both controls were considered putative binding proteins.

As we expected, HuR was highly enriched (>49 fold) in the HDAC2-TAG affinity purification, compared to the Control-TAG affinity purification. Thirteen additional proteins were enriched by HDAC2-TAG, which can be categorized as proteins involved in mRNA stabilization and splicing (Supplementary Data). In the 7SK-TAG affinity purification, we observed enrichment of canonical 7SK binding partners, including HEXIM1, CDK9, DDX21, MEPCE and LARP7 (Figure 3C). Other known binding partners, such as members of the BAF complex, were also identified.^34^ These results establish that our RNA-TAG approach can successfully identify RNA-binding proteins that are associated with specific RNAs.

### Application of RNA-TAG to target HOTAIR through minimal mutations

Use of RNA-TAG requires that a TGT recognition element, typically 17-25 nucleotides, be encoded into an RNA of interest in order to facilitate its labeling.^27^ Previous work has demonstrated that the minimal requirements for TGT recognition include a 7-nucleotide loop with the consensus sequence YUGUNNN, where Y represents a pyrimidine (C or U) and N represents any base.^36,37^ Furthermore, it has been hypothesized that recognition of an RNA substrate by TGT through charge complementarity requires adoption of a zig-zag conformation of the loop region.^38^ Because many RNAs, especially long noncoding RNAs, are highly structured,^39^ we next sought to explore whether existing stem loop structures within an RNA of interest could be leveraged to enable RNA-TAG labeling. HOTAIR, a 2,148 nt lncRNA, is overexpressed in several cancer tissues and has been implicated in various oncogenic processes including cell invasion, tumor development, and metastasis.^40^ Structural studies of HOTAIR have found it to be highly structured, with over 50% of the nucleotides involved in base-pairing interactions; furthermore, 56 helical segments were identified, with 38 terminal loops.^41^ We hypothesized that perhaps some of these existing stem-loops could be mutated in order to transform them into potential substrates for TGT. Therefore, we selected four candidate stem loops within HOTAIR that had a loop region of 7 nucleotides, H12, H22, H23, and H51 (Figure 4A). The loop regions of each of these stem loops were mutated to match the loop region of the TGT recognition element. These mutated HOTAIR transcripts were subjected to TGT labeling conditions, alongside a wild-type HOTAIR transcript, and control transcripts bearing a full TGT recognition element. Detection of biotinylation via northern blot showed that mutation of hairpin H22 (HOTAIR-H22-TAG) enabled labeling efficiencies similar to 5’TAG-HOTAIR, a transcript encoding a full 25-nucleotide TGT recognition element (Figure 4B, Supplementary Figure S3). This result was confirmed using labeling with preQ_1_-BODIPY and gel analysis (Supplementary Figure S4). Importantly, wild type HOTAIR transcript showed very low levels of detectable biotinylation. Despite extensive studies on the biological function of HOTAIR, its role in chromatin remodeling is still unclear and is a subject of debate.^42–46^ Therefore, we sought to employ our modified HOTAIR construct, along with the quantitative proteomics approach described above, in order to identify putative HOTAIR-binding proteins using MCF7-cells, a breast-cancer cell line known to express HOTAIR at high levels.^47,48^ Biotinylated HOTAIR-H22-TAG and 5’TAG-HOTAIR were compared to unlabeled controls, as well as a biotinylated transcript encoding a 5’TAG sequence and the antisense sequence corresponding to HOTAIR (5’TAG-antisense-HOTAIR), across five biological replicates. Biotinylation of the RNA constructs was verified by northern blot (Supplementary Figure S5). To generate lists of putative interacting proteins, the experimental sample (biotinylated target RNA) was compared to both controls (unlabeled target RNA and biotinylated 5’TAG-antisense-HOTAIR RNA). Proteins detected in at least three replicates with an average enrichment greater than 3-fold against both controls were considered putative binding proteins. Comparison of the samples enriched by the differentially labeled transcripts, 5’TAG-HOTAIR and HOTAIR-H22-TAG, identified 55 proteins that were enriched by both constructs. The ability to label HOTAIR with a minimal mutation allows for reduced perturbation of the RNA transcript. As such, we expected to observe a lower number of enriched proteins with the HOTAIR-H22-TAG RNA compared to the 5’TAG-HOTAIR due to fewer non-specific interactions. Several proteins involved in RNA processing, ribonucleoprotein biogenesis, and ribosome biogenesis were identified as binding proteins for both HOTAIR-TAG RNAs (Figure 4C, D, Supplementary Data).^49^ Interestingly, some of these proteins have been previously reported to have RNA-dependent functions in cancer. The long-noncoding RNA GSEC was previously shown to inhibit DHX36 through its g-quadruplexes, leading to cancer cell migration;^50^ importantly, an alternative g-quadruplex structure of HOTAIR has been reported.^51^ Furthermore, the oncogenic lncRNA MALAT1 has been proposed to influence alternative splicing through interactions with serine-arginine (SR) proteins, such as SRSF2.^52,53^. It has also been proposed that the long-noncoding RNA SLERT plays a role in the dysregulation of ribosome biogenesis in cancer, which may indicate a potential pathway for HOTAIR’s oncogenic function.^54^ While further studies are needed to characterize these binding partners, these findings suggest that HOTAIR may play alternative roles in oncogenesis that go beyond regulation of chromatin remodeling complexes.

**Figure 4.**
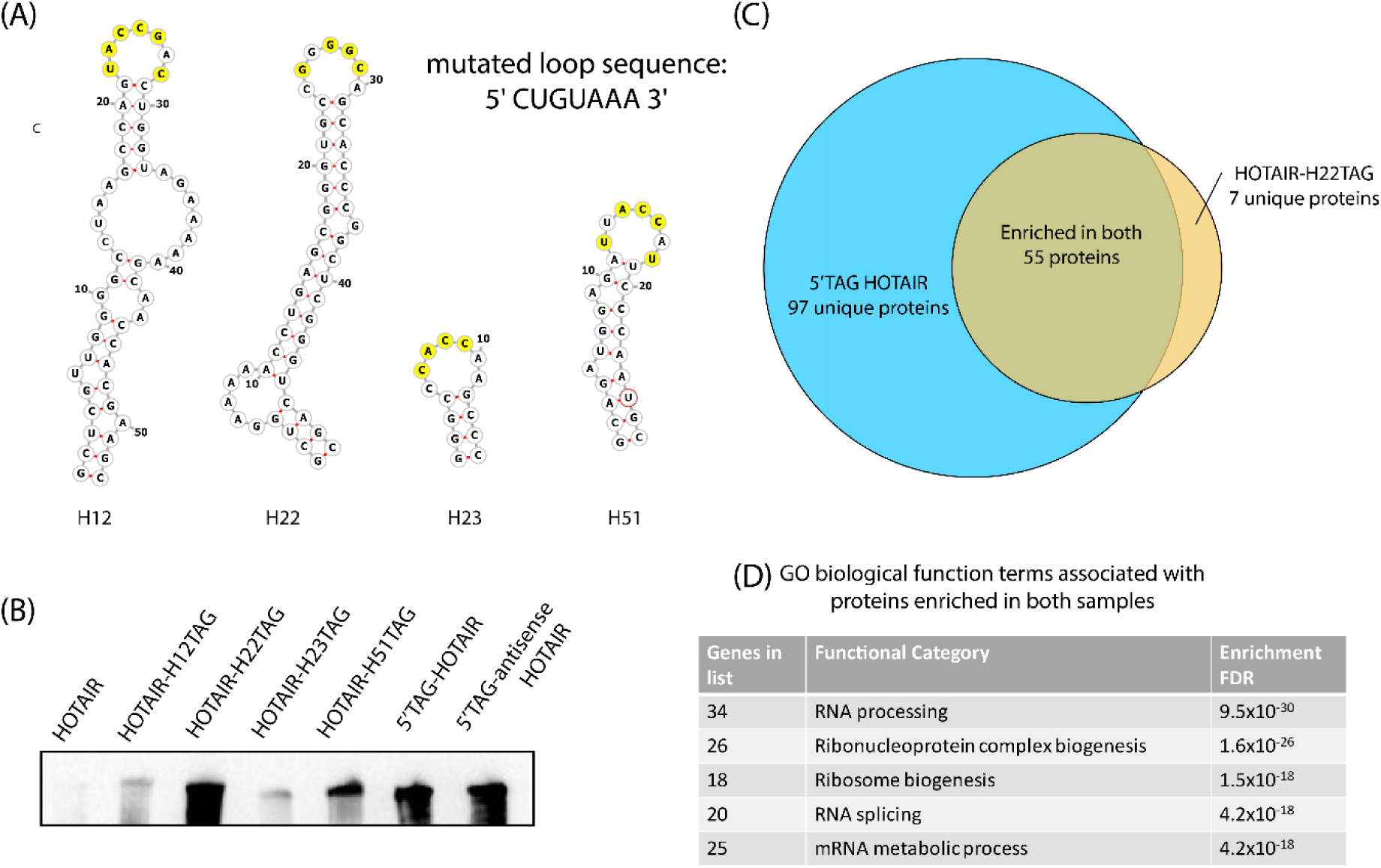
Analysis of HOTAIR-associated proteins with alternative labeling strategies. (A) Stem loop structures derived from previous structural characterization^41^ with yellow residues indicating bases that required mutation to match the loop (CUGUAAA) of the TGT recognition element. (B) Northern blot analysis of RNA transcripts subjected to TGT biotinylation conditions, developed with Streptavidin-HRP. (C) Venn diagram showing proteins that were enriched from MCF7 cell lysates >3 fold compared to both unlabeled and 5’TAG antisense-HOTAIR controls. (D) Top gene ontology biological function terms enriched in both samples.^49^

### Enrichment of cellular RNA with RNA-TAG

Having demonstrated the successful purification of various RNA-protein complexes using RNA-TAG labeling, we sought to determine, as a proof-of-principle, whether the RNA-TAG labeling approach was suitable for the purification of cellular RNA directly from cell lysates. In principle, this could allow for the study of cellular RNA-protein or RNA-RNA interactions. Using viral transduction, we prepared HeLa cells stably expressing the HDAC2-TAG RNA transcript. We reasoned that addition of TGT enzyme and preQ_1_-biotin directly to the corresponding cell lysate may enable selective biotinylation of the expressed transcript, which would allow for subsequent affinity purification with streptavidin-coupled beads. In this context, the selectivity of the TGT enzyme to label the targeted RNA over other RNAs present in the cell would be essential. It has been hypothesized that bacterial TGT forms a homodimer, where the first unit is catalytic and the second unit plays a role in recognition and proper orientation of the bound RNA substrate.^55^ Therefore, we postulated that a stable TGT dimer may have improved labeling selectivity and prepared an obligate TGT dimer, connected by a 16 amino acid XTEN linker, to test this hypothesis.^56^

We first sought to determine whether the expressed RNA could be efficiently and selectively labeled within a cellular lysate. TGT enzyme variants and preQ_1_-biotin were added to cell lysates from HeLa cells with stable HDAC2-TAG RNA expression, allowed to react, and total RNA was purified from the cell lysate to allow for RNA quantification. Streptavidin-mediated affinity purification was performed, and on-bead reverse transcription, followed by qPCR, was utilized to quantify affinity-purified, biotinylated RNA transcripts. As a control, the RNA input to the streptavidin affinity purification was also analyzed by RT-qPCR, so that the fraction of RNA recovered from a sample could be calculated (Figure 5A). From this analysis, we were gratified to find that 75% of the target HDAC2-TAG transcript was recovered using the *E. Coli* TGT enzyme and 44% was recovered using the TGT obligate dimer (Figure 5B). For comparison, an antisense oligonucleotide approach demonstrated ∼70% recovery of the long noncoding RNA XIST using RT-qPCR.^7^ Other RNA transcripts were also quantified by qPCR in order to test the selectivity of this method. Two RNAs, GAPDH mRNA and LBR mRNA, were selected due to their sequences, which contain a potential UGU-bearing stem-loop structure (Supplementary Table 1), with GAPDH also being a very abundant mRNA. Other abundant RNAs, including β-Actin mRNA, U1-snRNA, and 18s-rRNA were also tested. From this analysis, significant selectivity was observed against these RNAs using *E. coli* TGT, with GAPDH having a 1% recovery, and other RNAs having recoveries of less than 0.2% (Figure 5B, Supplementary Tables 2 and 3). An improvement in selectivity was observed in samples labeled with the obligate TGT dimer, specifically with the GAPDH transcript (0.02% recovery).

**Figure 5.**
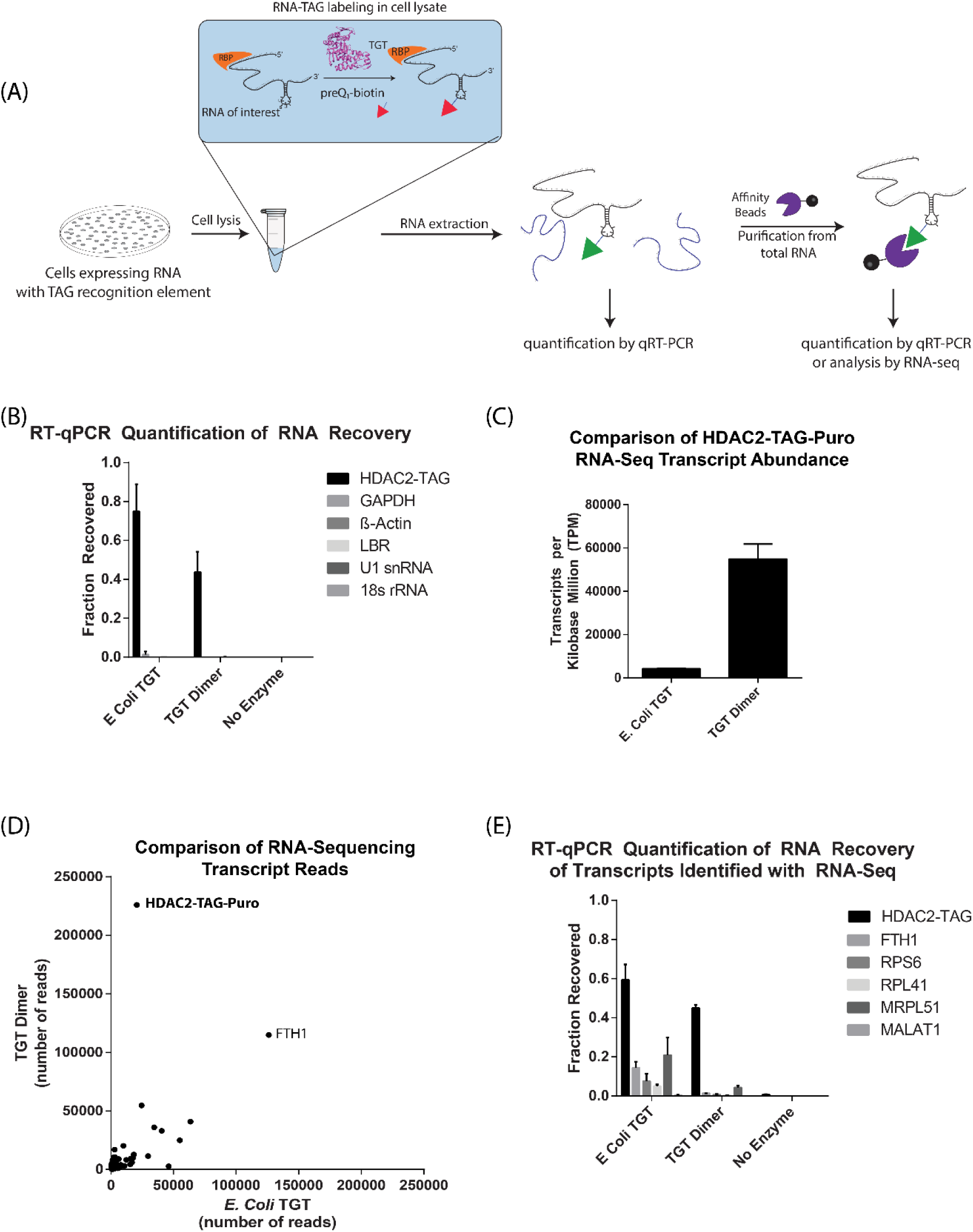
RNAs expressed in cells can be biotinylated efficiently and selectively in cell lysates. (A) Schematic of RNA labeling in cell lysate and RT-qPCR analysis. Cell lysates are supplemented with TGT enzyme and preQ_1_-biotin. Subsequently, RNA is extracted for affinity purification and analysis (B) Fraction of RNA recovered after cell lysate labeling was calculated (mean±S.D, N=3) by comparison of purified RNA to an input RNA sample, as described in methods. (C) Transcripts per kilobase million (TPM) of target HDAC2-TAG-Puro transcript in RNA sequencing libraries (mean±S.D, N=3). (D) Scatter plot comparing the average number of aligned reads for the top 1000 transcripts of the *E. Coli* TGT sample to TGT dimer sample. (E) Fraction of RNA recovered after cell lysate labeling was calculated for RNAs detected in the sequencing libraries (mean±S.D, N=3)

We then sought to further examine the selectivity of the monomeric and dimeric forms of TGT in the context of the human transcriptome through the preparation of RNA sequencing libraries. To accomplish this, we carried out lysate labeling, followed by streptavidin-mediated affinity purification of the extracted total RNA, and library preparation, using 3 replicates for each condition. Approximately 4-5 million reads were collected for each replicate, with reads aligning to greater than 45,000 RNA transcripts. Analysis of these libraries showed that the samples prepared with dimeric TGT had much more selective enrichment of the desired HDAC2-TAG-Puro transcript (Figures 5C, D). Furthermore, in each of the three replicates prepared with the obligate TGT dimer, the HDAC2-TAG-Puro transcript had the highest number of aligned reads compared to any other transcript, with an average of 10.95% of all aligned reads mapping to the HDAC2 construct. For comparison, RNA-sequencing was previously reported with the alternative RAP (RNA Antisense Purification) technique, with 68% of reads aligned to the targeted Xist transcript.^57^ Because the number of reads is dependent on transcript length, and Xist (∼17kb) is more than 8 times longer than the HDAC2-TAG-Puro transcript (∼2kb) studied here, this data is consistent with a similar degree of RNA selectivity amongst these techniques. Based on the observed improvement in selectivity for the obligate TGT dimer, we sought to further explore the selectivity of these two enzyme variants by further analysis of potential off-targets observed in the enriched sequencing libraries (FTH1, RPS6, RPL41, MRPL51, MALAT1). We examined whether these transcripts were enriched due to labeling by either monomeric or dimeric TGT through qPCR, as described above. This confirmed that the *E. coli* TGT enzyme displayed off-target labeling of several of these transcripts (5-21%), whereas the obligate TGT dimer off-target labeling was substantially reduced (Figure 5E, Supplementary Tables 4 and 5). This supports the hypothesis that the TGT homodimer aids in RNA recognition.

Based on the highly selective and efficient labeling of RNA with dimeric TGT, we hypothesized that this technique could be utilized to affinity purify an RNA of interest directly from cell lysates in the presence of remaining preQ_1_ biotin probe, which could potentially be applied to the affinity purification of cellular RNA-protein complexes. Treatment of cell lysates with enzymatic labeling conditions, followed directly with affinity purification and stringent washes allowed for direct enrichment of HDAC2-TAG RNA, approximately 130-fold, compared to a sample lacking the TGT enzyme. The obligate dimer showed similar levels of enrichment, approximately 145-fold, with significantly lower enrichment of the non-target GAPDH mRNA (Figure 6, Supplementary Tables 6 and 7). This data suggests that RNA labeling in cellular lysates using the obligate TGT dimer allows for selective, stringent purification of the expressed RNA, and could potentially be extended to the purification of cellular RNA-protein complexes directly from cell lysates.

**Figure 6.**
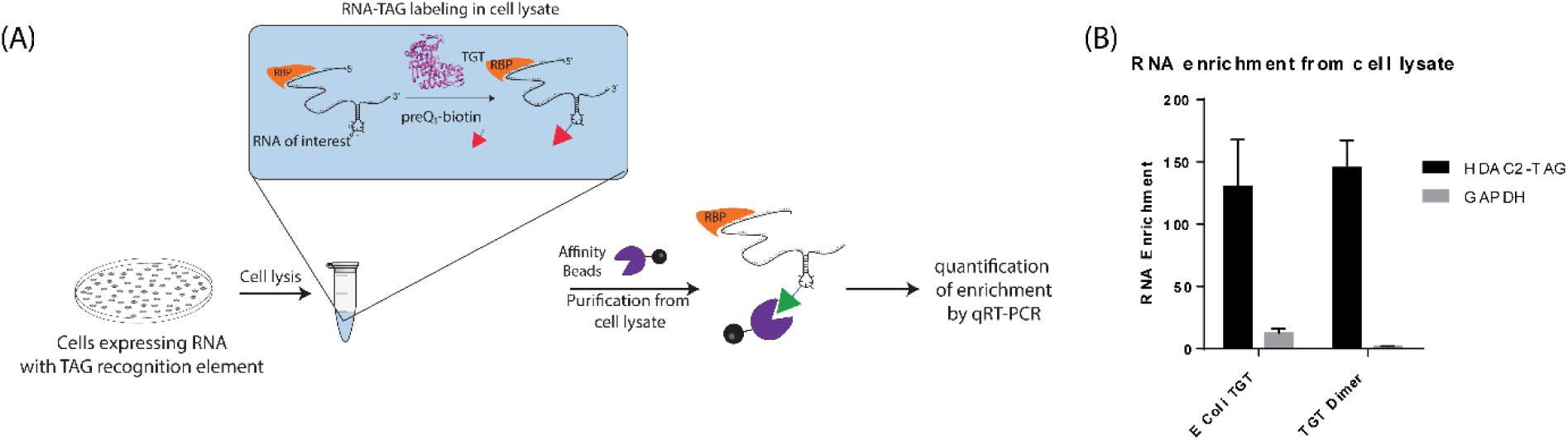
Labeling and purification of RNA directly from cell lysates. (A) RNA was labeled in cell lysates, and immediately purified using streptavidin beads and stringent washes, allowing for quantification of enrichment of RNA directly from cell lysates by RT-qPCR. (B) RT-qPCR enrichment as measured relative to no enzyme control, using β-actin as a reference gene, representing mean ± S.D., N=3. Supporting data can be found in the Supplementary Information.

### Conclusion

RNA-protein interactions regulate a broad range of critical cellular processes; however, identifying the proteins that interact with an RNA of interest in mammalian cells represents a major challenge in biology. While RNA-centric interaction mapping has been increasingly employed to elucidate RNA function, the noncovalent nature of existing approaches limits their utility to highly expressed RNA transcripts and represents a barrier to entry to many researchers due to the specialized and complex techniques employed. Direct covalent modification of an RNA of interest with biotin has many potential advantages for functional characterization. Due to the strength of the streptavidin-biotin interaction, RNA can be affinity purified when present at low concentrations and using stringent wash conditions. In this work, we present RNA-TAG as a novel methodology for the purification of RNA-protein complexes. By encoding a 25-nucleotide hairpin into mRNA constructs, analysis of RNA-protein complexes using immunoblotting was demonstrated using very low RNA concentrations. Furthermore, by combining the RNA-TAG methodology with quantitative proteomics, the identification of several known binding partners of the model 7SK snRNA was achieved.

In many cases, the addition of exogenous sequence elements to enable RNA labeling and isolation may be disadvantageous. In order to overcome this challenge, we mutated HOTAIR, a highly structured long noncoding RNA, to transform naturally occurring RNA hairpins into substrates for the TGT enzyme. Using this approach, the labeling of HOTAIR through the mutation of 4 nucleobases was utilized to enable proteomic analysis. This identified novel putative binding partners involved in pathways such as RNA processing, ribonucleoprotein biogenesis, RNA splicing, and ribosome biogenesis that may play a role in HOTAIR’s oncogenic functions.

Lastly, the ability of the RNA-TAG system to selectively label RNA expressed in mammalian cells was explored. Unlike other techniques for RNA enzymatic labeling, which do not enable targeted biotinylation of a specific transcript in a complex mixture,^21^ we demonstrated that RNA can be labeled selectively with good efficiency in cellular lysates, potentially allowing for this technique to be applied on a larger scale to isolate cellular RNA-protein complexes. While the bacterially expressed *E. Coli* TGT was found to be fairly promiscuous, engineering of a TGT dimer significantly improved RNA labeling selectivity, as demonstrated through RNA sequencing and RT-qPCR. Bacterial TGTs are thought to form a homodimer in cells, where one subunit is catalytic and the other subunit plays a role in binding and orienting the RNA substrate properly for catalysis;^55^ this observation of improved selectivity through the use of dimeric TGT supports this hypothesis. Using the dimeric TGT, the expressed RNA was labeled in cell lysates with good efficiency (44%), with selectivity similar to competitive affinity purification methods. For comparison, the use of tiling antisense oligonucleotides to purify the cellular lncRNA XIST enabled about ∼70% RNA recovery^7^ and similar selectivity by RNA sequencing,^57^ as discussed above. This approach is unique in its ability to efficiently and selectively biotinylate an RNA of interest in the presence of the human transcriptome, thus expanding the potential to affinity purify low abundance RNA transcripts.

Together, this report demonstrates a simple and highly flexible platform for the analysis of RNA-protein interactions through direct enzymatic biotinylation of a target RNA. This approach has several advantages, including the selectivity of RNA labeling against the human transcriptome, the ability to reduce the perturbation of the RNA recognition element through the mutation of existing structural elements, and the use of covalent biotinylation to achieve a strong and stringent affinity purification. The simplicity and robustness of the RNA-TAG approach creates opportunities for widespread utilization in studying RNA biology; we anticipate that this technology will aid in providing new insights into the functional characterization of RNA and its broad range of roles in cell biology.

## METHODS

### Synthesis of preQ_1_-biotin

PreQ_1_-C_6_H_12_-NH_2_ (2-amino-5-(((6-aminohexyl)amino)methyl)-3,7-dihydro-4H-pyrrolo[2,3-d]pyrimidin-4-one), prepared as previously described^58^, (2.5 mg, 9.0 µmol) was dissolved in 200 µL anhydrous DMF, followed by the addition of anhydrous diisopropylethylamine (4.6 µL, 27 µmol). A solution of biotin-NHS ester (3.2 mg, 9.3 µmol) in 200 µL DMF was then slowly added to the mixture. The reaction was stirred for 2 h at room temperature. The reaction was concentrated, and the residue was purified by HPLC to give a white solid (2.2 mg, 49% yield).

### In vitro transcription

All plasmids used for in vitro transcription were prepared using standard cloning techniques. The plasmid encoding Control-TAG is available on Addgene (Addgene #138209, pcDNA3.1- (empty)-TAG). This vector backbone was used to prepare HDAC2-TAG, β-actin-TAG, and 7SK-TAG. The plasmid encoding HOTAIR was provided by Anna Pyle^41^. T7 RNA polymerase was expressed and purified as previously reported^26^. Templates were linearized via restriction digest for Control-TAG (XbaI), HDAC2-TAG(XbaI), β-actin-TAG (XbaI) and HOTAIR constructs (SalI). PCR amplification was used to linearize 7SK-TAG and append a T7 promoter with the forward primer AAGCTGTAATACGACTCACTATAGGGGATCCCCGGGAGCAGAC and the reverse primer AAAAGAAAGGCAGACTGCCACATG. Transcription reactions were set up with RNA NTPs (5 mM each ATP, CTP, UTP, 9 mM GTP) (NEB, Ipswitch, MA), 0.004 U/µL Thermostable Inorganic Pyrophosphatase (NEB, Ipswitch, MA), 0.15 μg/μL T7-RNAP, and 0.05% Triton X-100 (Sigma, St. Louis, MO) in T7 Reaction Buffer. Each transcription reaction was setup with approximately 4 µg of cut plasmid or 1 µg of purified PCR product as a template in a 100 μL transcription reaction. The transcription reaction was run at 37 °C for 3-4 hours. RNA was then purified via lithium chloride precipitation by addition of LiCl Precipitation Solution (Invitrogen, Carlsbad, CA) to reach a final concentration of 2.5 M LiCl, followed by incubation at -20 °C for 1 h or overnight. The RNA was quantified at 260 nm, confirmed as a single observable UV shadowing band by 4% denaturing PAGE (4% polyacrylamide in TBE with 8M urea) and kept frozen at -20 °C until used.

### TGT labeling of RNA Transcripts

*E. Coli* tRNA Guanine Transglycosylase (TGT) was expressed and purified as previously described^26^. TGT labeling reactions were carried out in 1X TGT reaction buffer with 5 mM DTT, 1 µM RNA transcript, 1 µM *E. Coli* TGT, 10 µM preQ_1_-biotin, and 1 U/µL Murine RNAse Inhibitor (NEB, Ipswitch, MA). Labeling reactions were incubated for 2 h at 37 °C, and purified via ethanol precipitation.

### Northern Blot Biotinylation Assay

Northern blot biotinylation assays were carried out with 2 pmol of RNA for HDAC2-TAG, 7SK-TAG, and Control-TAG transcripts, and 0.1 pmol of RNA for all HOTAIR constructs. RNA was electrophoretically separated by 4% denaturing PAGE (4% polyacrylamide in TBE with 8M urea), stained with SYBR green II (Invitrogen, Carlsbad, CA) and imaged. Subsequently, RNA was transferred via electroblot to a positively charged nylon membrane (Invitrogen, Carlsbad, CA) in 0.5X TBE. Blot was developed for biotin detection using the Chemiluminescent Nucleic Acid Detection Module Kit (Thermo Scientific, Waltham, MA)

### Cell Culture

Human HeLa-S3 cells were grown in complete DMEM media (Gibco) supplemented with 10% fetal bovine serum and 1% penicillin/streptomycin. Human MCF7 HTB-22 cells were grown in Eagle’s MEM (Gibco) supplemented with 10% fetal bovine serum, 1mM pyruvate, 0.01 mg/ml human recombinant insulin and 1% penicillin/streptomycin. Cells were cultured at 37°C in a humidified incubator under 5% CO_2_.

### RNA-Protein Analysis by Immunoblot

Biotinylated RNA was folded in 1X RNA folding buffer (10 mM Tris pH 7, 0.1 M KCl, 10 mM MgCl_2_) by heating for 2 min at 90 °C, followed by 20 min at 25 °C. HeLa cell pellets (∼3×10^6^) were lysed with 500 µL Mammalian Protein Extraction Reagent (Thermo Scientific), supplemented with 2X HALT protease inhibitor (Thermo Scientific), according to manufacturer’s instructions. Protein concentration was quantified using Pierce BCA Protein Assay Kit (Thermo Scientific). Folded RNA (1 pmol, unless otherwise noted) was incubated with 40 µg HeLa cell lysate in 1X Immunoblot Binding and Washing Buffer (1X IBW, 25 mM Tris-HCl pH 7.5, 150 mM KCl, 2 mM MgCl_2_, 1 mM DTT, 0.25% Nonidet-P40, 1% Tween 20), supplemented with 1 U/µL Murine RNAse Inhibitor (NEB, Ipswitch, MA) for 1 h at room temperature to allow binding of cognate proteins. 10 µL washed Dynabeads M-280 Streptavidin (Invitrogen, Carlsbad, CA) were added to each binding reaction and further incubated at RT for one hour. Beads were washed 3 times in 1X IBW Buffer, followed by 2 washes with PBS. Protein was eluted from beads by heating for 5 min at 95 °C in 1X Laemmli Buffer (Bio-Rad, Hercules, CA). Retrieved protein was detected by standard immunoblotting techniques using the following antibodies: mouse anti-HuR (3A2, Santa Cruz Biotechnology) and mouse anti-β-tubulin (D-10, Santa Cruz Biotechnology).

### RNA-Protein Enrichment, Trypsinization, and Stable Isotope Labeling for Mass Spectrometry

HeLa cell pellets (∼2×10^7^) or MCF7 cell pellets (∼4×10^7^) were lysed with 3 mL Mammalian Protein Extraction Reagent (Thermo Scientific), supplemented with 2X HALT protease inhibitor (Thermo Scientific), according to manufacturer’s instructions. Protein concentration was quantified using Pierce BCA Protein Assay Kit (Thermo Scientific). Biotinylated RNAs were labeled as described above, while unlabeled RNA controls were used directly. RNA was folded in 1X RNA folding buffer by heating for 2 min at 90 °C, followed by 20 min at 25 °C. For HDAC2-TAG and 7SK-TAG experiments, each folded RNA (5 pmol) was incubated with 200 µg HeLa cell lysate in 1X Proteomics Binding and Washing Buffer (1X PBW, 20 mM Potassium Phosphate pH 7.5, 150 mM KCl, 2 mM MgCl_2_, 1 mM DTT, 0.25% Nonidet P40, 1% Tween 20), supplemented with 1 U/µL Murine RNAse Inhibitor (NEB, Ipswitch, MA) for 1 h at room temperature to allow binding of cognate proteins. For HOTAIR experiments, each folded RNA (10 pmol) was incubated with 400 µg MCF7 cell lysate in 1X PBW, supplemented with 1 U/µL Murine RNAse Inhibitor (NEB, Ipswitch, MA) for 1 h at room temperature to allow binding of cognate proteins. 25 µL washed Dynabeads M-280 Streptavidin (Invitrogen, Carlsbad, CA) were added to each binding reaction and further incubated at RT for one hour. Beads were washed 3 times in 1X PBW Buffer, followed by 5 washes with PBS. Beads were then resuspended in 50 µL trypsinization solution consisting of 10 ng/µL Trypsin (Promega) in 100 mM tetramethylammonium bicarbonate (TEAB), pH 8.0. Trypsinization was carried out overnight at 37 °C with sample rotation. The supernatant containing trypsinized peptides was then transferred to a new tube and vacuum dried. Stable isotope dimethyl labeling was carried out according to previously published protocols^35^. The samples were labeled as indicated in the Supplementary Data.

### Mass Spectrometry

The labeled, vacuum dried peptides were resuspended in 5% Formic acid/5% Acetonitrile buffer and added to the vials for mass spectrometry analysis. Samples were analyzed with triplicate injections by LC-MS-MS using EASY-nLC 1000 liquid chromatography connected with Q-Exactive mass spectrometer (Thermo Scientific, San Jose, CA) as described previously^59^ with some modification as follows. The peptides were eluted using the 60 minute Acetonitrile gradient (45 minutes 2%-30% ACN gradient followed by 5 minutes 30%-60% ACN gradient, a 2 minute 60-95% ACN gradient, and a final 8-minute isocratic column equilibration step at 0% ACN) at 250nL/minute flow rate. All the gradient mobile phases contained 0.1% formic acid. The data dependent analysis (DDA) was done using top 10 method with a positive polarity, scan range 400-1800 m/z, 70,000 resolution, and an AGC target of 3e6. A dynamic exclusion time of 20 s was implemented and unassigned, singly charged and charge states above 6 were excluded for the data dependent MS/MS scans. The MS2 scans were triggered with a minimum AGC target threshold of 1e5 and with maximum injection time of 60 ms. The peptides were fragmented using the normalized collision energy (NCE) setting of 25. Apex trigger and peptide match settings were disabled.

### Mass Spectrometry Analysis

RAW files were processed, searched, and analyzed essentially as described previously.^60^ Converted mzXML files were searched using SEQUEST (version 28) using a target-decoy database containing reviewed UniProtKB/Swiss-Prot Human protein sequences and common contaminants. Each mzXML file was searched in triplicate with the following parameters: 50 parts per million precursor ion tolerance and 0.01-Da fragment ion tolerance; Trypsin (1 1 KR P) was set as the enzyme; up to three missed cleavages were allowed; a dynamic modification of 15.99491 Da on methionine (oxidation); and a static modification of 57.02146 Da on cysteine for iodoacetamide alkylation. For searches with light and medium reductive dimethyl labels, additional dynamic modifications of 4.0224 Da on lysine and peptide N termini and static modifications of 28.0313 Da on lysine and peptide N termini were included. For searches with light and heavy reductive dimethyl labels, additional dynamic modifications of 8.04437 Da on lysine and peptide N termini and static modifications of 28.0313 Da on lysine and peptide N termini were included. For searches with medium and heavy reductive dimethyl labels, additional dynamic modifications of 4.02193 Da on lysine and peptide N termini and static modifications of 32.05374 Da on lysine and peptide N termini were included. Peptide matches were filtered to a peptide false discovery rate of 2% using the linear discriminant analysis. Proteins were further filtered to a false discovery rate of 2%, peptides were assembled into proteins using maximum parsimony, and only unique and razor peptides were retained for subsequent analysis. All peptide heavy/light, medium/light, and heavy/medium ratios with a signal-to-noise ratio of above 5 were used for assembled protein quantitative ratios.

### Generation of cell lines stably expressing HDAC2-TAG

Stable cell lines expressing HDAC2-TAG was generated using lentiviral transduction and subsequent puromycin selection as follows. 293T cells were transfected with the following helper plasmids pHAGE - GAG-POL; pHAGE - VSVG; pHAGE - tat1b; pHAGE - rev and pHAGE-HDAC2-TAG-IRES-Puro using Mirus TransIT 293 transfection reagent. After 24 h fresh media was added to the cells. 48 hour post transfection, the media containing the infectious virions was collected and filtered using a 0.45 mm sterile syringe filter. Polybrene (6ug/mL) was added to the filtered virus-containing media. The mixture was added to HeLa cells seeded at 30% confluency and infected for 24hours, prior to selection with 500 ng/mL Puromycin to obtain stable expression clones.

### Expression and purification of obligate dimeric TGT

A plasmid encoding obligate dimeric TGT was cloned from the TGT-His plasmid (Addgene #138201) using DNA HiFi Assembly (New England Biolabs) to add a 16-amino acid XTEN linker (SGSETPGTSESATPES) between two identical coding sequences for *E. Coli* TGT. Enzyme was then expressed in BL21-DE3 cells with pG-KJE8 chaperone plasmid (Takara Bio) in media supplemented with 0.5 mg/mL arabinose and 5 ng/mL doxycycline to induce chaperone expression, and 1 mM IPTG to induce enzyme expression. Expression was allowed to proceed overnight at 18 °C, and dimeric TGT was then purified using standard His-tag purification, as described previously^26^.

### RNA Labeling in cell lysate

Stable HeLa cells expressing HDAC2-TAG, grown to ∼80% confluency in a 10 cm plate, were lysed directly upon addition of 600 µL of Mammalian Protein Extraction Reagent (Thermo Scientific), supplemented with 2X HALT protease inhibitor (Thermo Scientific) and 1 U/µL Murine RNAse, according to manufacturer’s instructions. To this cell lysate, 1 µM preQ_1_-biotin, 100 nM *E. Coli* TGT or 500 nM obligate dimeric TGT (as noted), and 1 U/µL Murine RNAse inhibitor was added. Reactions were incubated for 2 h at room temperature prior to analysis.

### qPCR analysis of RNA Recovery

Following RNA labeling in cell lysate, as described above, total RNA was extracted from the lysate upon the addition of 3 volumes of Trizol-LS (Invitrogen) and purification with the Zymo Direct-Zol kit (Zymo Scientific) according to manufacturer’s instructions. 10 µL of washed Dynabeads MyOne C1 Streptavidin (Invitrogen) were incubated with 200 ng isolated total RNA sample in 1X RNA Binding and Washing Buffer (1X RBW, 5 mM Tris-HCl, pH 7.5, 0.5 mM EDTA, 1 M NaCl, 0.05% Tween) supplemented with 2 U/µL Murine RNAse inhibitor for 1 h at room temperature. Beads were washed 3 times in 1X RBW Buffer, followed by 2 washes with ultra-pure water. On-bead reverse transcription was performed in a volume of 20 μl using 200 U of Maxima Reverse Transcriptase (Thermo Scientific) according to the product instructions. For input controls, 200 ng of the corresponding total RNA sample was used. Following reverse transcription, cDNA was diluted 6-fold, and qPCR was performed using 4 μl of diluted cDNA template in a 20 μl reaction volume using qPCRBIO SyGreen Blue Mix (PCR Biosystems). Quantitative PCR was performed on a CFX-96 Real-Time System with a C100 Thermocycler (Bio-Rad). RNA recovery values were determined by comparison of the input and recovered RNA C_T_ values by computing the ΔC_T_(purified-input) according to the following equation: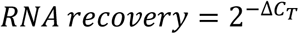, based on the assumptions of the ΔΔC_T_ method.^61^ Primer sequences are available in the Supplementary Information.

### Enrichment of labeled RNA from cell lysate

Following RNA labeling in cell lysate, as described above, 25 µL of washed Dynabeads MyOne C1 Streptavidin (Invitrogen) was added directly to the lysate labeling reaction and allowed to incubate for 1 h at room temperature. Beads were washed 3 times in 1X Stringent Wash Buffer I (20 mM Tris-HCl, pH 8, 2 mM EDTA, 150 mM NaCl, 1% Triton-X, 0.1% SDS), followed by 3 washes with 1X Stringent Wash Buffer II (20 mM Tris-HCl, pH 8, 2 mM EDTA, 500 mM NaCl, 1% Triton-X, 0.1% SDS), and finally 2 washes with ultra-pure water. On-bead reverse transcription was performed in a volume of 20 μl using 200 U of Maxima Reverse Transcriptase (Thermo Scientific) and a cocktail of 15 pmol each of gene-specific RT primers, according to the product instructions. Following on-bead reverse transcription, we performed qPCR and determined the ΔC_T_(gene-reference) for each condition, using β-Actin as a reference. We then calculated the ΔΔC_T_ between the enzyme treated sample and control sample lacking enzyme treatment, and then calculated enrichment according to the following equation: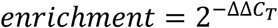, based on the assumptions of the ΔΔC_T_ method.^61^

### RNA Sequencing

In order to identify transcripts biotinylated using this method, RNA was biotinylated in cell lysate, as described above. Total RNA was extracted from the lysate upon the addition of 3 volumes of Trizol-LS (Invitrogen) and purification with the Zymo Direct-Zol kit (Zymo Scientific) according to manufacturer’s instructions. Library preparation was conducted using TruSeq Stranded mRNA kit (Illumina), using a modified protocol. In lieu of polyA purification, 25 µL of washed Dynabeads MyOne C1 Streptavidin (Invitrogen) were incubated with 10 µg isolated total RNA sample in 1X RBW Buffer supplemented with 2 U/µL Murine RNAse inhibitor for 1 h at room temperature. Beads were washed 3 times in 1X RBW Buffer, followed by 2 washes with ultra-pure water. Subsequent to RNA binding and washing to the streptavidin beads, 19.5 μL of Fragment-Prime-Finish mix was added to each beads sample to elute and fragment the RNA. Library preparation was carried out in subsequent steps according to manufacturer’s protocol. All libraries were then sequenced using a MiSeq (Illumina) to produce paired end 75-bp reads.

### RNA sequencing raw data processing

Raw RNA-seq data was first separated in paired fragment mates. Sequence AGATCGGAAGAGCACACGTCTGAACTCCAGTCA or AGATCGGAAGAGCGTCGTGTAGGGAAAGAGTGT was used to cut potential sequencing adaptor reads from raw RNA-seq read_1 or read_2, respectively. Sequencing reads were aligned using Bowtie2 in end-to-end mapping mode with index built from the human hg38 assembly and the transgene sequence. Salmon^62^ was used to map RNA-seq reads to human transcriptome and quantify transcript expression.

## Supporting information

Supplementary Data - MS Hitlists

## AVAILABILITY

Plasmids for TGT enzyme (#138201) and Control-TAG RNA (#138209) are available on Addgene. RNA sequencing data has been deposited to GEO with accession number GSE149274.

## ASSOCIATED CONTENT

The Supporting Information is available on BioRxiv.

## ACKNOWLEDGEMENT

The authors gratefully acknowledge Anna Pyle for providing the plasmid for transcription of HOTAIR and UCSD Molecular Mass Spectrometry Facility for small molecule high resolution mass spectrometry support. This work was supported by the National Institutes of Health [R01 GM123285 to N.K.D. and E.J.B., T32 GM112584-03 to K.N.B.].

## Notes

https://www.ncbi.nlm.nih.gov/geo/query/acc.cgi?acc=GSE149274

### Competing Interest Statement

The authors have declared no competing interest.

