## Supplementary Information for "Enzymatic RNA Biotinylation for Affinity Purification and Identification of RNA-protein Interactions"

#### Supplementary Figures and Tables

##### Supplementary Figure S1: Quantification of RNA recovery with reduced RNA concentrations

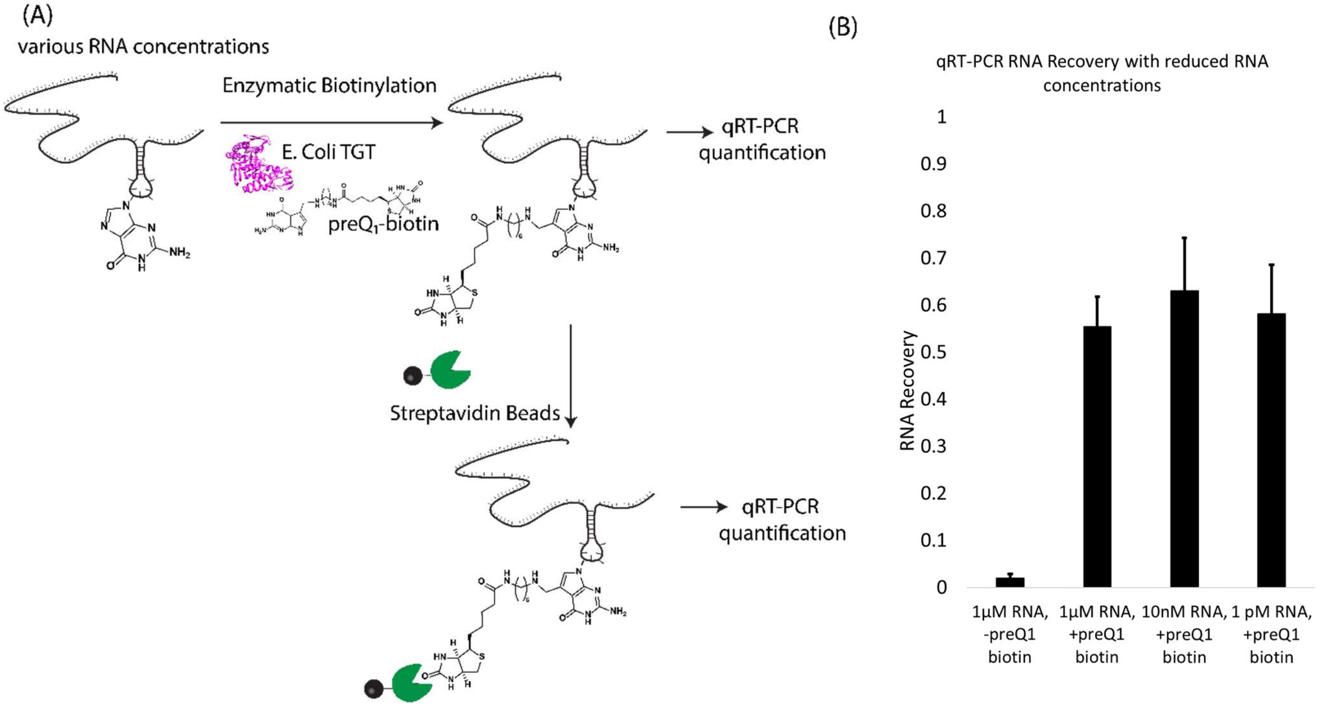


(A)HDAC2-TAG RNA at the indicated concentrations (1µM, 10 nM, 1 pM) was treated with preQ$_1$-biotin and TGT under standard labeling conditions. After precipitation of the RNA product, RNA was diluted to uniform concentrations for RT-qPCR analysis. Input RNA was quantified by qRT-PCR, alongside RNA that was affinity purified using Dynabeads M-280 Streptavidin. (B) RNA recovery values were determined by comparison of the input and recovered RNA C$_T$ values, as described in Methods. Biotinylation of RNA was observed in RNA concentrations as low as 1 pM.

#### Supplementary Figure S2: Northern blot assay of HDAC2-TAG and 7SK-TAG RNA transcripts used in mass spectrometry experiments

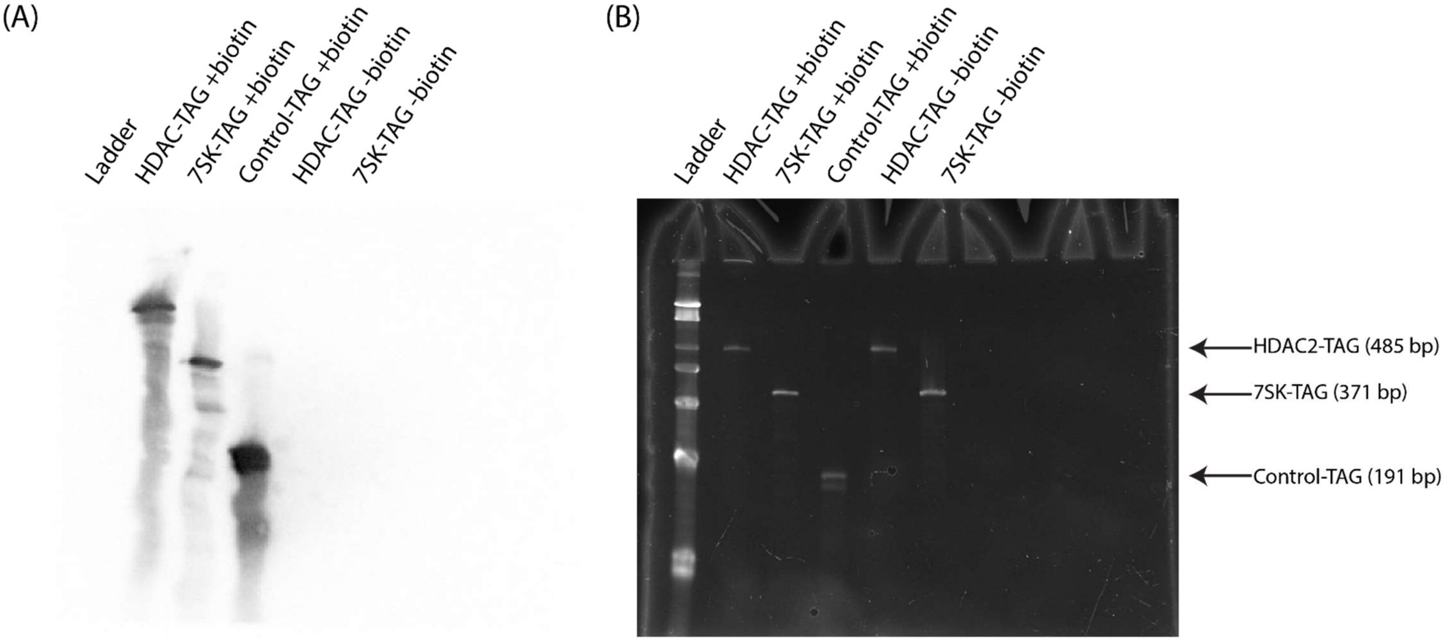


Northern blot results for transcripts used in HDAC2-TAG and 7SK-TAG proteomics experiments (A) Biotin detection (B) SYBR green staining.

#### Supplementary Figure S3: Northern blot assay of HOTAIR mutant labeling

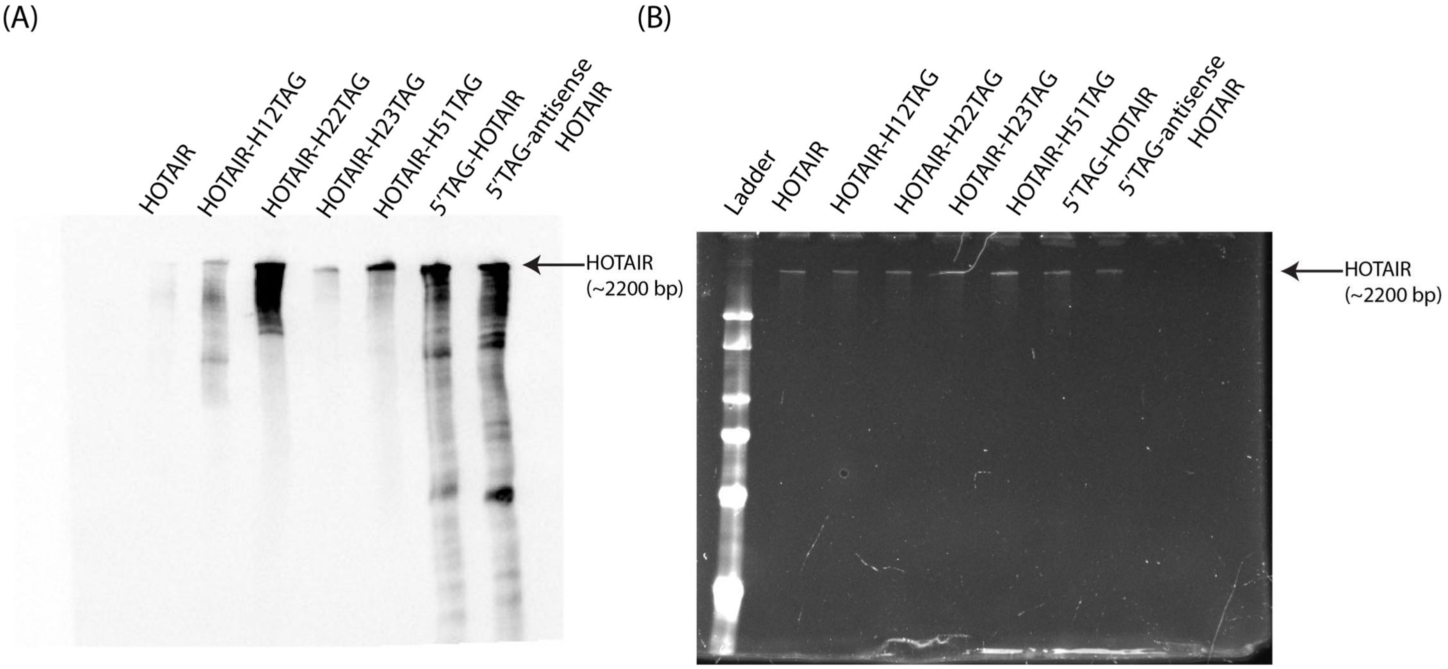


Northern blot results of HOTAIR labeling assay (A) Biotin detection (B) SYBR green staining.

#### Supplementary Figure S4: HOTAIR labeling with preQ1-BODIPY

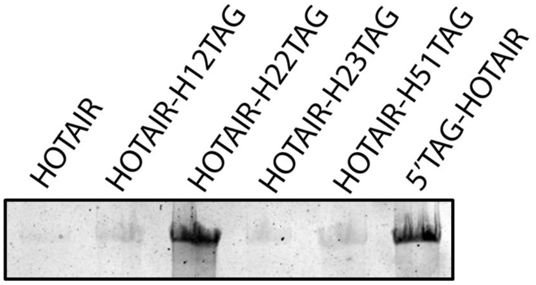


Polyacrylamide gel electrophoresis of HOTAIR constructs treated with TGT and preQ$_1$-BODIPY. 1µM RNA was treated with 1 µM *E. Coli* TGT and 10 µM preQ$_1$-BODIPY[1] for 2 h at 37°C. RNA was subsequently precipitated with lithium chloride and analyzed via 4% denaturing UREA-PAGE. BODIPY fluorescence was detected using a Bio-Rad ChemiDoc-MP imager.

#### Supplementary Figure S5: Northern blot assay of HOTAIR transcripts used in mass spectrometry experiments

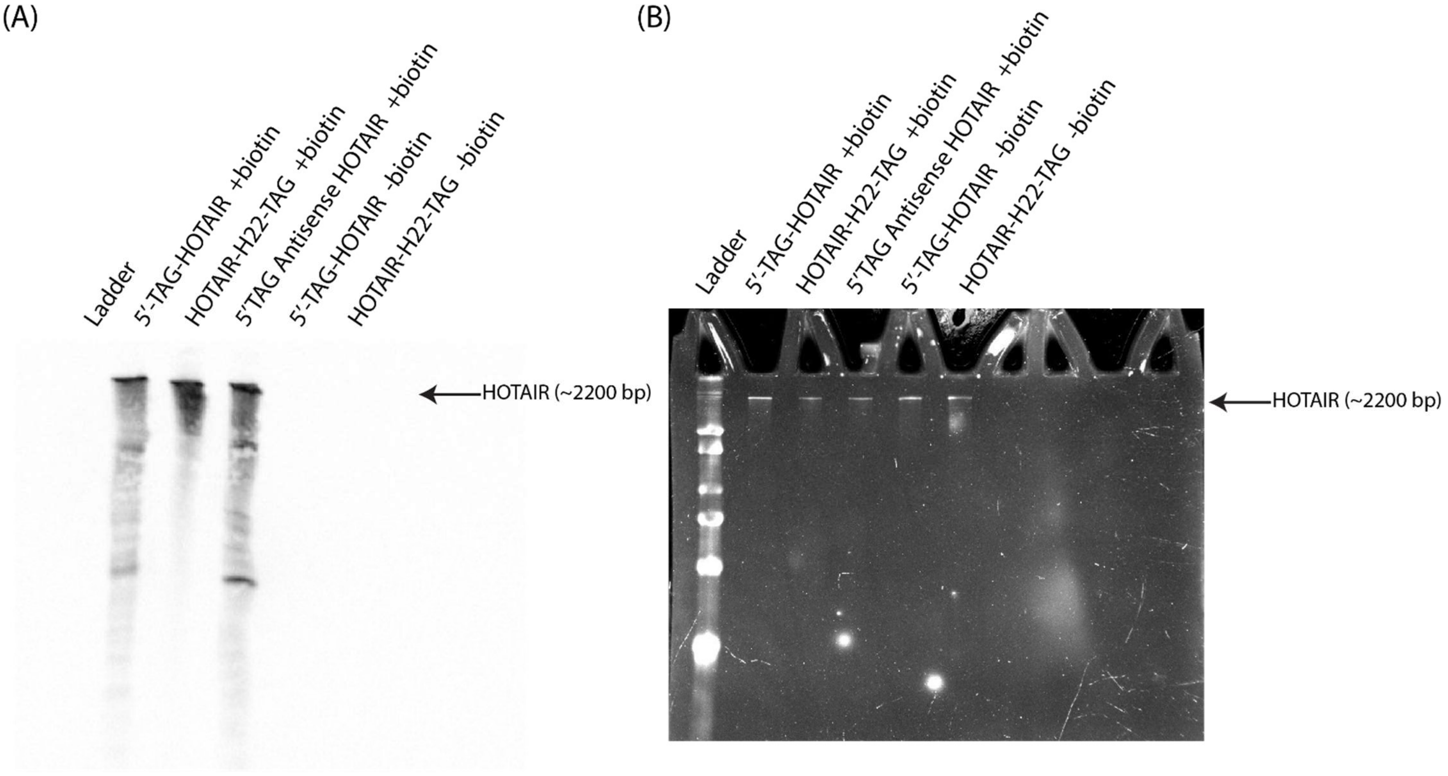


Northern blot results for transcripts used in HOTAIR proteomics experiments (A) Biotin detection (B) SYBR green staining.

#### Supplementary Table 1: Putative TAG-like sequences in human transcripts

| Gene | Transcript accession number | TAG-like cDNA Sequence |
|---|---|---|
| GAPDH | NM_001256799.3 | 'CATGG**GTGTGAA**CCATG' |
| LBR | NM_002296.4 | 'ACTCC**CTGTAAA**GGAGT' |
| MRPL51 | NM_016497.4 | 'GGAAA**CTGTACT**TTTCC' |
| MALAT1 | NR_002819.4 | 'AGAAA**ATGTTTT**TTTCT' |

#### Supplementary Table 2: qPCR ΔCt data of initial selectivity screen

| | | ΔCt (purified-input) | | |
|---|---|---|---|---|
| | | E Coli TGT | TGT Dimer | No Enzyme |
| **HDAC2-TAG** | Rep1 | 0.34 | 0.85 | 13.03 |
| | Rep2 | 0.21 | 1.5 | 15.29 |
| | Rep3 | 0.75 | 1.31 | 16.98 |
| **GAPDH** | Rep1 | 5.00 | 12.83 | 15.19 |
| | Rep2 | 8.01 | 11.8 | 21.24 |
| | Rep3 | 8.57 | 11.72 | 19.17 |
| **Actin** | Rep1 | 10.13 | 16.45 | 17.03 |
| | Rep2 | 14.24 | 13.83 | 18.18 |
| | Rep3 | 12.5 | 17.54 | 17.16 |
| **LBR** | Rep1 | 9.68 | 10.46 | 12.03 |
| | Rep2 | 9.76 | 10.3 | 12.96 |
| | Rep3 | 9.47 | 13.25 | 13.67 |
| **U1 snRNA** | Rep1 | 16.8 | 20.65 | 13.89 |
| | Rep2 | 19.09 | 21.17 | 20.15 |
| | Rep3 | 17.61 | 15.14 | 12.72 |
| **18s rRNA** | Rep1 | 11.37 | 18.2 | 11.6 |
| | Rep2 | 13.56 | 17.82 | 18.81 |
| | Rep3 | 13.85 | 14.95 | 10.61 |

#### Supplementary Table 3: Calculated qPCR Recovery of initial selectivity screen

| | Calculated RNA Recovery (mean ± S.D., 3 replicates) | | |
|---|---|---|---|
| | E Coli TGT | TGT Dimer | No Enzyme |
| **HDAC2-TAG** | 75.0 ± 13.9% | 43.7 ± 10.5% | 0.0051 ± 0.0060% |
| **GAPDH** | 1.25 ± 1.62% | 0.024 ± 0.009% | 0.0010 ± 0.0015% |
| **Actin** | 0.037 ± 0.045% | 0.0028 ± 0.0035% | 0.0006 ± 0.0002% |
| **LBR** | 0.13 ± 0.01% | 0.054 ± 0.038% | 0.015 ± 0.008% |
| **U1 snRNA** | 0.0005 ± 0.0004% | 0.0010 ± 0.0016% | 0.0072 ± 0.0074% |
| **18s rRNA** | 0.018 ± 0.018% | 0.0013 ± 0.0016% | 0.032 ± 0.032% |

#### Supplementary Table 4: qPCR ΔCt data examining RNA-Seq Transcripts

| | | dCt (purified-input) | | |
|---|---|---|---|---|
| | | E Coli TGT | TGT Dimer | No Enzyme |
| **HDAC2-TAG** | Rep1 | 0.82 | 1.14 | 7.08 |
| | Rep2 | 0.55 | 1.11 | 7.38 |
| | Rep3 | 0.91 | 1.21 | 6.88 |
| **FTH1** | Rep1 | 2.98 | 6.21 | 12.62 |
| | Rep2 | 2.47 | 6.26 | 12.59 |
| | Rep3 | 2.98 | 6.15 | 12.38 |
| **RPS6** | Rep1 | 4.27 | 7.18 | 13.37 |
| | Rep2 | 3.07 | 6.72 | 13.7 |
| | Rep3 | 4.13 | 7.11 | 12.93 |
| **RPL41** | Rep1 | 4.27 | 8.07 | 14.68 |
| | Rep2 | 4.11 | 8.04 | 15.06 |
| | Rep3 | 4.61 | 8.25 | 15.34 |
| **MRPL51** | Rep1 | 2.65 | 4.75 | 10.34 |
| | Rep2 | 1.68 | 4.22 | 11.1 |
| | Rep3 | 2.66 | 4.74 | 11.33 |
| **MALAT1** | Rep1 | 8.71 | 12.36 | 15.57 |
| | Rep2 | 7.32 | 11.55 | 15.89 |
| | Rep3 | 8.95 | 12.16 | 16.22 |

#### Supplementary Table 5: Calculated qPCR Recovery Data of RNA-Seq Transcripts

| | Calculated RNA Recovery (mean ± S.D., 3 replicates) | | |
|---|---|---|---|
| | E Coli TGT | TGT Dimer | No Enzyme |
| **HDAC2-TAG** | 59.4 ± 7.9 % | 45.0 ± 1.6% | 0.73 ± 0.12% |
| **FTH1** | 14.5 ± 3.1% | 1.35 ± 0.05% | 0.017 ± 0.002% |
| **RPS6** | 7.60 ± 3.74% | 0.79 ± 0.14% | 0.010 ± 0.003% |
| **RPL41** | 5.02 ± 0.86% | 0.36 ± 0.03% | 0.003 ± 0.001% |
| **MRPL51** | 21.0 ± 8.9% | 4.27 ± 0.95 % | 0.054 ± 0.021% |
| **MALAT1** | 0.36 ± 0.23% | 0.025 ± 0.008% | 0.0017 ± 0.0004% |

#### Supplementary Table 6: qPCR Ct values from lysate purification

| | | Rep1 | Rep2 | Rep3 |
|---|---|---|---|---|
| **HDAC2-TAG** | E Coli TGT | 19.97304 | 19.4716 | 19.509 |
| | TGT dimer | 20.42991 | 19.9254 | 20.206 |
| | No enzyme | 27.73386 | 26.1111 | 27.77 |
| **GAPDH** | E Coli TGT | 21.16293 | 20.2132 | 20.522 |
| | TGT dimer | 24.16379 | 23.8605 | 23.997 |
| | No enzyme | 25.1854 | 24.3525 | 25.03 |
| **Actin** | E Coli TGT | 25.03013 | 24.2149 | 24.018 |
| | TGT dimer | 25.30845 | 25.0633 | 25.064 |
| | No enzyme | 25.39997 | 24.3181 | 25.256 |

#### Supplementary Table 7: Calculated Enrichment values from lysate purification

| | E Coli TGT | TGT dimer |
|---|---|---|
| **HDAC** | 130.2 ± 37.5 | 145.3 ± 22.0 |
| **GAPDH** | 12.9 ± 3.4 | 2.0 ± 0.3 |

### Supplementary Notes

#### Gene Sequences

The gene sequences for each construct are listed below (from the T7 promoter to the restriction cut site, or transcription stop site), with the TAG recognition element underlined. Mutated HOTAIR constructs were prepared using Q5 site directed mutagenesis with the following primers:

| **Primer Name** | Sequence |
|---|---|
| HOTAIR-H12TAG-Fwd | aaaCTGGTAGAAAAAGCAACCACGAAG |
| HOTAIR-H12TAG-Rev | acagCTGGCTTAGGCCCCAACG |
| HOTAIR-H22TAG-Fwd | aaaGCACCCGGCTCGGGTCAG |
| HOTAIR-H22TAG-Rev | acagGCACCCGCTCAGGTTTTTCCAG |
| HOTAIR-H23TAG-Fwd | aaaGCCCCGCCCCTCGCGGCC |
| HOTAIR-H23TAG-Rev | acagGCCCGGTGTGGGGCAGTGGC |
| HOTAIR-H51TAG-Fwd | aaaTCCCAATGCCTGAACTTC |
| HOTAIR-H51TAG-Rev | acagTCTCCATCTGCTGATTTTTTTTC |

*Control-TAG*

GGGAGACCCAAGCTGGCTAGCGTTTAAACTTAAGCTTGGTACCGAGCTCGGATCCTCTAACACAGACTCTCGGTACCATCATTTTCAT
ATCCCCGGGAGCAGACTGTAAATCTGCTCCCACCACCATCATTTAATGAATTCCATCAGGAATCCCTCACTTCTGCAGACTGGCCGTCG
TTTTACACTCGAGT

*HDAC2-TAG*

GGGAGACCCAAGCTGGCTAGCGTTTAAACTTAAGCTTGGTACCGAGCTCGGATCCTGACAGTCTCACCAATTTCAGAAAATCATTAAA
AAGAAAATATTGAAAGGAAAATGTTTTCTTTTTGAAGACTTCTGGCTTCATTTTATACTACTTTGGCATGGACTGTATTTATTTTCAAAT
GGCTTTTTCGTTTTTGTTTTTCTTGGCAAGTTTTATTGTGAGTTTTTCTAATTATGAAGCAAAATTTCTTTTCTCCACCATGCTTTATGTG
ATAGTATTTAAAATTGATGTGAGTTATTATGTCAAAAAAACTGATCTATTAAAGAAGTAATTGGCCTTTCTGAGCTGATATCTAACACA
GACTCTCGGTACCATCATTTTCATATCCCCGGGAGCAGACTGTAAATCTGCTCCCACCACCATCATTTAATGAATTCCATCAGGAATCC
CTCACTTCTGCAGACTGGCCGTCGTTTTACACTCGAGT

*β-Actin-TAG*

GGGAGACCCAAGCTGGCTAGCACCGCCGAGACCGCGTCCGCCCCGCGAGCACAGAGCCTCGCCTTTGCCGATCCGCCGCCCGTCCAC
ACCCGCCGCCAGCTCACCATGGATGATGATATCGCCGCGCTCGTCGTCGACAACGGCTCCGGCATGTGCAAGGCCGGCTTCGCGGGC
GACGATGCCCCCCGGGCCGTCTTCCCCTCCATCGTGGGGCGCCCCAGGCACCAGGGCGTGATGGTGGGCATGGGTCAGAAGGATTC
CTATGTGGGCGACGAGGCCCAGAGCAAGAGAGGCATCCTCACCCTGAAGTACCCCATCGAGCACGGCATCGTCACCAACTGGGACG
ACATGGAGAAAATCTGGCACCACACCTTCTACAATGAGCTGCGTGTGGCTCCCGAGGAGCACCCCGTGCTGCTGACCGAGGCCCCCC
TGAACCCCAAGGCCAACCGCGAGAAGATGACCCAGATCATGTTTGAGACCTTCAACACCCCAGCCATGTACGTTGCTATCCAGGCTGT
GCTATCCCTGTACGCCTCTGGCCGTACCACTGGCATCGTGATGGACTCCGGTGACGGGGTCACCCACACTGTGCCCATCTACGAGGG
GTATGCCCTCCCCCATGCCATCCTGCGTCTGGACCTGGCTGGCCGGGACCTGACTGACTACCTCATGAAGATCCTCACCGAGCGCGGC
TACAGCTTCACCACCACGGCCGAGCGGGAAATCGTGCGTGACATTAAGGAGAAGCTGTGCTACGTCGCCCTGGACTTCGAGCAAGA
GATGGCCACGGCTGCTTCCAGCTCCTCCCTGGAGAAGAGCTACGAGCTGCCTGACGGCCAGGTCATCACCATTGGCAATGAGCGGTT
CCGCTGCCCTGAGGCACTCTTCCAGCCTTCCTTCCTGGGCATGGAGTCCTGTGGCATCCACGAAACTACCTTCAACTCCATCATGAAGT
GTGACGTGGACATCCGCAAAGACCTGTACGCCAACACAGTGCTGTCTGGCGGCACCACCATGTACCCTGGCATTGCCGACAGGATGC
AGAAGGAGATCACTGCCCTGGCACCCAGCACAATGAAGATCAAGATCATTGCTCCTCCTGAGCGCAAGTACTCCGTGTGGATCGGCG
GCTCCATCCTGGCCTCGCTGTCCACCTTCCAGCAGATGTGGATCAGCAAGCAGGAGTATGACGAGTCCGGCCCCTCCATCGTCCACCG
CAAATGCTTCTAGGCGGACTATGACTTAGTTGCGTTACACCCTTTCTTGACAAAACCTAACTTGCGCAGAAAACAAGATGAGATTGGC
ATGGCTTTATTTGTTTTTTTTGTTTTGTTTTGGTTTTTTTTTTTTTTTTGGCTTGACTCAGGATTTAAAAACTGGAACGGTGAAGGTGACA
GCAGTCGGTTGGAGCGAGCATCCCCCAAAGTTCACAATGTGGCCGAGGACTTTGATTGCACATTGTTGTTTTTTTAATAGTCATTCCA
AATATGAGATGCGTTGTTACAGGAAGTCCCTTGCCATCCTAAAAGCCACCCCACTTCTCTCTAAGGAGAATGGCCCAGTCCTCTCCCA
AGTCCACACAGGGGAGGTGATAGCATTGCTTTCGTGTAAATTATGTAATGCAAAATTTTTTTAATCTTCGCCTTAATACTTTTTTATTTT
GTTTTATTTTGAATGATGAGCCTTCGTGCCCCCCCTTCCCCCTTTTTTGTCCCCCAACTTGAGATGTATGAAGGCTTTTGGTCTCCCTGG
GAGTGGGTGGAGGCAGCCAGGGCTTACCTGTACACTGACTTGAGACCAGTTGGGATCCTCTAACACAGACTCTCGGTACCATCATTT
TCATATCCCCGGGAGCAGACTGTAAATCTGCTCCCACCACCATCATTTAATGAATTCCATCAGGAATCCCTCACTTCTGCAGACTGGCC
GTCGTTTTACACTCGAGT

*7SK-TAG*

GGATCCCCGGGAGCAGACTGTAAATCTGCTCCCACCACCGGATGTGAGGGCGATCTGGCTGCGACATCTGTCACCCCATTGATCGCC
AGGGTTGATTCGGCTGATCTGGCTGGCTAGGCGGGTGTCCCCTTCCTCCCTCACCGCTCCATGTGCGTCCCTCCCGAAGCTGCGCGCT
CGGTCGAAGAGGACGACCATCCCCGATAGAGGAGGACCGGTCTTCGGTCAAGGGTATACGAGTAGCTGCGCTCCCCTGCTAGAACC
TCCAAACAAGCTCTCAAGGTCCATTTGTAGGAGAACGTAGGGTAGTCAAGCTTCCAAGACTCCAGACACATCCAAATGAGGCGCTGC
ATGTGGCAGTCTGCCTTTCTTTT

*HOTAIR*

GGGACTCGCCTGTGCTCTGGAGCTTGATCCGAAAGCTTCCACAGTGAGGACTGCTCCGTGGGGGTAAGAGAGCACCAGGCACTGAG
GCCTGGGAGTTCCACAGACCAACACCCCTGCTCCTGGCGGCTCCCACCCGGGGCTTAGACCCTCAGGTCCCTAATATCCCGGAGGTG
CTCTCAATCAGAAAGGTCCTGCTCCGCTTCGCAGTGGAATGGAACGGATTTAGAAGCCTGCAGTAGGGGAGTGGGGAGTGGAGAG

AGGGAGCCCAGAGTTACAGACGGCGGCGAGAGGAAGGAGGGGCGTCTTTATTTTTTTAAGGCCCCAAAGAGTCTGATGTTTACAAG
ACCAGAAATGCCACGGCCGCGTCCTGGCAGAGAAAAGGCTGAAATGGAGGACCGGCGCCTTCCTTATAAGTATGCACATTGGCGAG
AGAATTAAGTGCTGCAACCTAAACCAGCAATTACACCCAAGCTCGTTGGGGCCTAAGCCAGTACCGACCTGGTAGAAAAAGCAACCA
CGAAGCTAGAGAGAGAGCCAGAGGAGGGAAGAGAGCGCCAGACGAAGGTGAAAGCGAACCACGCAGAGAAATGCAGGCAAGGG
AGCAAGGCGGCAGTTCCCGGAACAAACGTGGCAGAGGGCAAGACGGGCACTCACAGACAGAGGTTTATGTATTTTTATTTTTTAAAA
TCTGATTTGGTGTTCCATGAGGAAAAGGGAAAATCTAGGGAACGGGAGTACAGAGAGAATAATCCGGGTCCTAGCTCGCCACATGA
ACGCCCAGAGAACGCTGGAAAAACCTGAGCGGGTGCCGGGGCAGCACCCGGCTCGGGTCAGCCACTGCCCCACACCGGGCCCACCA
AGCCCCGCCCCTCGCGGCCACCGGGGCTTCCTTGCTCTTCTTATCATCTCCATCTTTATGATGAGGCTTGTTAACAAGACCAGAGAGCT
GGCCAAGCACCTCTATCTCAGCCGCGCCCGCTCAGCCGAGCAGCGGTCGGTGGGGGGACTGGGAGGCGCTAATTAATTGATTCCTTT
GGACTGTAAAATATGGCGGCGTCTACACGGAACCCATGGACTCATAAACAATATATCTGTTGGGCGTGAGTGCACTGTCTCTCAAAT
AATTTTTCCATAGGCAAATGTCAGAGGGTTCTGGATTTTTAGTTGCTAAGGAAAGATCCAAATGGGACCAATTTTAGGAGGCCCAAA
CAGAGTCCGTTCAGTGTCAGAAAATGCTTCCCCAAAGGGTTGGCAGTGTGTTTTGTTGGAAAAAAGCTTGGGTTATAGGAAAGCCTT
TCCCTGCTACTTGTGTAGACCCAGCCCAATTTAAGAATTACAAGGAAGCGAAGGGGTTGTGTAGGCCGGAAGCCTCTCTGTCCCGGC
TGGATGCAGGGGACTTGAGCTGCTCCGGAATTTGAGAGGAACATAGAAGCAAAGGTCCAGCCTTTGCTTCGTGCTGATTCCTAGACT
TAAGATTCAAAAACAAATTTTTAAAAGTGAAACCAGCCCTAGCCTTTGGAAGCTCTTGAAGGTTCAGCACCCACCCAGGAATCCACCT
GCCTGTTACACGCCTCTCCAAGACACAGTGGCACCGCTTTTCTAACTGGCAGCACAGAGCAACTCTATAATATGCTTATATTAGGTCTA
GAAGAATGCATCTTGAGACACATGGGTAACCTAATTATATAATGCTTGTTCCATACAGGAGTGATTATGCAGTGGGACCCTGCTGCA
AACGGGACTTTGCACTCTAAATATAGGCCCCAGCTTGGGACAAAAGTTGCAGTAGAAAAATAGACATAGGAGAACACTTAAATAAGT
GATGCATGTAGACACAGAAGGGGTATTTAAAAGACAGAAATAATAGAAGTACAGAAGAACAGAAAAAAAATCAGCAGATGGAGAT
TACCATTCCCAATGCCTGAACTTCCTCCTGCTATTAAGATTGCTAGAGAATTGTGTCTTAAACAGTTCATGAACCCAGAAGAACGCAAT
TTCAATGTATTTAGTACACACACAGTATGTATATAAACACAACTCACAGAATATATTTCCATACATTGGGTAGGTATGCACTTTGTGT
ATATATAATAATGTATTTTCCATGCAGTTTTAAAATGTAGATATATTAATATCTGGATGCATTTTCG

*5’TAG-HOTAIR*

GGGGGAGCAGACTGTAAATCTGCTCCCGACTCGCCTGTGCTCTGGAGCTTGATCCGAAAGCTTCCACAGTGAGGACTGCTCCGTGGG
GGTAAGAGAGCACCAGGCACTGAGGCCTGGGAGTTCCACAGACCAACACCCCTGCTCCTGGCGGCTCCCACCCGGGGCTTAGACCC
TCAGGTCCCTAATATCCCGGAGGTGCTCTCAATCAGAAAGGTCCTGCTCCGCTTCGCAGTGGAATGGAACGGATTTAGAAGCCTGCA
GTAGGGGAGTGGGGAGTGGAGAGAGGGAGCCCAGAGTTACAGACGGCGGCGAGAGGAAGGAGGGGCGTCTTTATTTTTTTAAGG
CCCCAAAGAGTCTGATGTTTACAAGACCAGAAATGCCACGGCCGCGTCCTGGCAGAGAAAAGGCTGAAATGGAGGACCGGCGCCTT
CCTTATAAGTATGCACATTGGCGAGAGAATTAAGTGCTGCAACCTAAACCAGCAATTACACCCAAGCTCGTTGGGGCCTAAGCCAGT
ACCGACCTGGTAGAAAAAGCAACCACGAAGCTAGAGAGAGAGCCAGAGGAGGGAAGAGAGCGCCAGACGAAGGTGAAAGCGAAC
CACGCAGAGAAATGCAGGCAAGGGAGCAAGGCGGCAGTTCCCGGAACAAACGTGGCAGAGGGCAAGACGGGCACTCACAGACAG
AGGTTTATGTATTTTTATTTTTTAAAATCTGATTTGGTGTTCCATGAGGAAAAGGGAAAATCTAGGGAACGGGAGTACAGAGAGAAT
AATCCGGGTCCTAGCTCGCCACATGAACGCCCAGAGAACGCTGGAAAAACCTGAGCGGGTGCCGGGGCAGCACCCGGCTCGGGTCA
GCCACTGCCCCACACCGGGCCCACCAAGCCCCGCCCCTCGCGGCCACCGGGGCTTCCTTGCTCTTCTTATCATCTCCATCTTTATGATG
AGGCTTGTTAACAAGACCAGAGAGCTGGCCAAGCACCTCTATCTCAGCCGCGCCCGCTCAGCCGAGCAGCGGTCGGTGGGGGGACT
GGGAGGCGCTAATTAATTGATTCCTTTGGACTGTAAAATATGGCGGCGTCTACACGGAACCCATGGACTCATAAACAATATATCTGTT
GGGCGTGAGTGCACTGTCTCTCAAATAATTTTTCCATAGGCAAATGTCAGAGGGTTCTGGATTTTTAGTTGCTAAGGAAAGATCCAAA
TGGGACCAATTTTAGGAGGCCCAAACAGAGTCCGTTCAGTGTCAGAAAATGCTTCCCCAAAGGGTTGGCAGTGTGTTTTGTTGGAAA
AAAGCTTGGGTTATAGGAAAGCCTTTCCCTGCTACTTGTGTAGACCCAGCCCAATTTAAGAATTACAAGGAAGCGAAGGGGTTGTGT
AGGCCGGAAGCCTCTCTGTCCCGGCTGGATGCAGGGGACTTGAGCTGCTCCGGAATTTGAGAGGAACATAGAAGCAAAGGTCCAGC
CTTTGCTTCGTGCTGATTCCTAGACTTAAGATTCAAAAACAAATTTTTAAAAGTGAAACCAGCCCTAGCCTTTGGAAGCTCTTGAAGGT
TCAGCACCCACCCAGGAATCCACCTGCCTGTTACACGCCTCTCCAAGACACAGTGGCACCGCTTTTCTAACTGGCAGCACAGAGCAAC
TCTATAATATGCTTATATTAGGTCTAGAAGAATGCATCTTGAGACACATGGGTAACCTAATTATATAATGCTTGTTCCATACAGGAGTG
ATTATGCAGTGGGACCCTGCTGCAAACGGGACTTTGCACTCTAAATATAGGCCCCAGCTTGGGACAAAAGTTGCAGTAGAAAAATAG
ACATAGGAGAACACTTAAATAAGTGATGCATGTAGACACAGAAGGGGTATTTAAAAGACAGAAATAATAGAAGTACAGAAGAACAG
AAAAAAAATCAGCAGATGGAGATTACCATTCCCAATGCCTGAACTTCCTCCTGCTATTAAGATTGCTAGAGAATTGTGTCTTAAACAG
TTCATGAACCCAGAAGAACGCAATTTCAATGTATTTAGTACACACACAGTATGTATATAAACACAACTCACAGAATATATTTTCCATAC

ATTGGGTAGGTATGCACTTTGTGTATATATAATAATGTATTTTCCATGCAGTTTTAAAATGTAGATATATTAATATCTGGATGCATTTTC
G

*5'TAG Antisense HOTAIR*

GGGGGAGCAGACTGTAAATCTGCTCCCGAAAATGCATCCAGATATTAATATATCTACATTTTAAAACTGCATGGAAAATACATTATTA
TATATACACAAAGTGCATACCTACCCAATGTATGGAAAATATATTCTGTGAGTTGTGTTTATATACATACTGTGTGTGTACTAAATACA
TTGAAATTGCGTTCTTCTGGGTTCATGAACTGTTTAAGACACAATTCTCTAGCAATCTTAATAGCAGGAGGAAGTTCAGGCATTGGGA
ATGGTAATCTCCATCTGCTGATTTTTTTTCTGTTCTTCTGTACTTCTATTATTTCTGTCTTTTAAATACCCCTTCTGTGTCTACATGCATCA
CTTATTTAAGTGTTCTCCTATGTCTATTTTTCTACTGCAACTTTTGTCCCAAGCTGGGGCCTATATTTAGAGTGCAAAGTCCCGTTTGCA
GCAGGGTCCCACTGCATAATCACTCCTGTATGGAACAAGCATTATATAATTAGGTTACCCATGTGTCTCAAGATGCATTCTTCTAGACC
TAATATAAGCATATTATAGAGTTGCTCTGTGCTGCCAGTTAGAAAAGCGGTGCCACTGTGTCTTGGAGAGGCGTGTAACAGGCAGGT
GGATTCCTGGGTGGGTGCTGAACCTTCAAGAGCTTCCAAAGGCTAGGGCTGGTTTCACTTTTAAAAATTTGTTTTTGAATCTTAAGTCT
AGGAATCAGCACGAAGCAAAGGCTGGACCTTTGCTTCTATGTTCCTCTCAAATTCCGGAGCAGCTCAAGTCCCCTGCATCCAGCCGGG
ACAGAGAGGCTTCCGGCCTACACAACCCCTTCGCTTCCTTGTAATTCTTAAATTGGGCTGGGTCTACACAAGTAGCAGGGAAAGGCTT
TCCTATAACCCAAGCTTTTTTCCAACAAAACACACTGCCAACCCTTTGGGGAAGCATTTTCTGACACTGAACGGACTCTGTTTGGGCCT
CCTAAAATTGGTCCCATTTGGATCTTTCCTTAGCAACTAAAAATCCAGAACCCTCTGACATTTGCCTATGGAAAAATTATTTGAGAGAC
AGTGCACTCACGCCCAACAGATATATTGTTTATGAGTCCATGGGTTCCGTGTAGACGCCGCCATATTTTACAGTCCAAAGGAATCAAT
TAATTAGCGCCTCCCAGTCCCCCCACCGACCGCTGCTCGGCTGAGCGGGCGCGGCTGAGATAGAGGTGCTTGGCCAGCTCTCTGGTC
TTGTTAACAAGCCTCATCATAAAGATGGAGATGATAAGAAGAGCAAGGAAGCCCCGGTGGCCGCGAGGGGCGGGGCTTGGTGGGC
CCGGTGTGGGGCAGTGGCTGACCCGAGCCGGGTGCTGCCCCGGCACCCGCTCAGGTTTTTCCAGCGTTCTCTGGGCGTTCATGTGGC
GAGCTAGGACCCGGATTATTCTCTCTGTACTCCCGTTCCCTAGATTTTCCCTTTTCCTCATGGAACACCAAATCAGATTTTAAAAAATAA
AAATACATAAACCTCTGTCTGTGAGTGCCCGTCTTGCCCTCTGCCACGTTTGTTCCGGGAACTGCCGCCTTGCTCCCTTGCCTGCATTTC
TCTGCGTGGTTCGCTTTCACCTTCGTCTGGCGCTCTCTTCCCTCCTCTGGCTCTCTCTCTAGCTTCGTGGTTGCTTTTTCTACCAGGTCGG
TACTGGCTTAGGCCCCAACGAGCTTGGGTGTAATTGCTGGTTTAGGTTGCAGCACTTAATTCTCTCGCCAATGTGCATACTTATAAGG
AAGGCGCCGGTCCTCCATTTCAGCCTTTTCTCTGCCAGGACGCGGCCGTGGCATTTCTGGTCTTGTAAACATCAGACTCTTTGGGGCC
TTAAAAAAATAAAGACGCCCCTCCTTCCTCTCGCCGCCGTCTGTAACTCTGGGCTCCCTCTCTCCACTCCCCACTCCCCTACTGCAGGCT
TCTAAATCCGTTCCATTCCACTGCGAAGCGGAGCAGGACCTTTCTGATTGAGAGCACCTCCGGGATATTAGGGACCTGAGGGTCTAA
GCCCCGGGTGGGAGCCGCCAGGAGCAGGGGTGTTGGTCTGTGGAACTCCCAGGCCTCAGTGCCTGGTGCTCTCTTACCCCCACGGA
GCAGTCCTCACTGTGGAAGCTTTCGGATCAAGCTCCAGAGCACAGGCGAGTCG

#### Primers used for RT-qPCR

| Primer | Sequence |
|---|---|
| HDAC2-TAG RT | CTCCCGGGGATATGAAAATG |
| GAPDH RT | GTGAAGACGCCAGTG |
| β-actin RT | GTGGATGCCACAGGAC |
| LBR RT | CTTCATAATAAAGTGAACTCCCAG |
| 18S RT | GAGGGCCTCACTAAACC |
| U1 RT | CCCACTACCACAAATTATGC |
| HDAC2-TAG qPCR fwd | AATTTCTTTTCTCCACCATGCTTTATGTG |
| HDAC2-TAG qPCR rev | GAAAATGATGGTACCGAGAGTCTGTGTTAG |
| GAPDH qPCR fwd | AATCCCATCACCATCTTCCA |
| GAPDH qPCR rev | TGGACTCCACGACGTACTCA |
| β-actin qPCR fwd | AGAGCTACGACGTGCCTGAC |
| β-actin qPCR rev | CTCCATGCCCAGGAAGGAAGG |
| LBR qPCR fwd | TGCTGTGCGACTATTCTCC |
| LBR qPCR rev | CAGGCCATCGACCTCTTACC |
| 18S qPCR fwd | GTAACCCGTTGAACCCC |
| 18S qPCR rev | CCATCCAATCGGTAGTAGCG |
| U1 qPCR fwd | CCATGATCACGAAGGTGGTTT |

| | |
|---|---|
| U1 qPCR rev | ATGCAGTCGAGTTTCCCACAT |
| FTH1 qPCR fwd | CCAGAACTACCACCAGGACTC |
| FTH1 qPCR rev | GTCAAAGTAGTAAGACATGGACAGG |
| RPS6 qPCR fwd | AAGCACCCAAGATTCAGCGT |
| RPS6 qPCR rev | TAGCCTCCTTCATTCTCTTGGC |
| RPL41 qPCR fwd | AGCCAAGTGGAGGAAGAAGC |
| RPL41 qPCR rev | AGCGTCTGGCATTCCATGTT |
| MALAT1 qPCR fwd | TGGTGATGAAGGTAGCAGGC |
| MALAT1 qPCR rev | ATTGCCGACCTCACGGATTT |
| MRPL51 qPCR fwd | AAGCTTCTCTCTTGGTGTGC |
| MRPL51 qPCR rev | CCAGGATCCCGATGTTGTCA |

#### $preQ_1$-biotin Characterization Data

$^1H$ NMR (500 MHz, $CD_3OD$) δ 6.85 (s, 1H), 4.50 (dd, *J* = 7.8, 4.4 Hz, 1H), 4.31 (dd, *J* = 7.9, 4.5 Hz, 1H), 3.24 – 3.15 (m, 4H), 3.06 (t, *J* = 9.5 Hz, 2H), 2.93 (dd, *J* = 12.6, 5.1 Hz, 1H), 2.71 (d, *J* = 12.7 Hz, 1H), 2.20 (t, *J* = 7.3 Hz, 3H), 1.79 – 1.32 (m, 18H). $^{13}C$ NMR (126 MHz, $CD_3OD$) δ 174.62, 166.64, 164.73, 161.25, 153.10, 152.57, 117.69, 108.43, 98.21, 61.98, 60.19, 55.66, 46.19, 43.37, 39.64, 38.57, 35.37, 28.76, 28.38, 28.12, 25.89, 25.62, 25.57.

HRMS [M+H]+ m/z calcd. for [C23H37N8O3S] + 505.2704, found 505.2707 (Δ = 0.6 ppm).
